# cGAS bends unpaired DNA to form an unconventional structure that hyperactivates the innate immune response

**DOI:** 10.64898/2026.05.22.726336

**Authors:** Shuangshuang Yang, Shuai Wu, Silian Chen, Xin Li, Iryna Chelepis, Smaranda Willcox, Katherine C. Barnett, Xiaoqing Hu, Guannan Huang, Willie J. Brickey, Joseph A Duncan, Wei-Chun Chou, Pengda Liu, William G Fusco, Stephanie M. Torres, Gerald S. Shadel, Gregory D. Bowman, Jack D. Griffith, Jungsan Sohn, Jenny P.-Y. Ting

## Abstract

cGAS is a pattern-recognition receptor for dsDNA and forms 2cGAS:2DNA dimers followed by oligomerization into phase-separated condensates when fully-complementary DNA is studied. However, many DNAs are not fully complementary. We report that DNA with unpaired regions such as those found during transcription, recombination or replication (designated as bubble-DNA, Bu-DNA) causes cGAS hyper-activation. Hyperactivation is observed by Bu-DNA embedded in linear DNA, circular DNA, plasmid DNA and mitochondria DNA. Bu-DNA binds significantly more tightly to the cGAS catalytic domain than paired-DNA but suppresses condensation. Cryo-EM and single-molecule FRET reveal that cGAS forms 2cGAS:1DNA complexes by bending Bu-DNA into a V-shape using the unpaired region as a hinge, limiting its oligomeric state. This uncovers a novel mode of cGAS activation attributed to pattern diversity within pattern ligands.

## Main text

In eukaryotic cells, genomic DNA is normally sequestered within the nucleus and mitochondria. However, under cellular stresses, such as genotoxic DNA damage, defective mitophagy, or pathogen invasion, DNA can aberrantly breach these organelles and enter the cytosol (*1*). To detect such abnormal signatures, known as pathogen-associated or damage-associated molecular patterns (PAMPs or DAMPs), the innate immune system has evolved an array of pattern recognition receptors (PRRs) that initiate inflammatory signaling cascades upon activation (*2, 3*). Over the past several decades, elucidating how cells sense cytosolic DNA has been pivotal to understanding host defenses against pathogens and mechanisms of cellular homeostasis and inflammatory pathology. Among DNA sensors, cyclic GMP-AMP synthase (cGAS) has emerged as a predominant PRR, with direct roles in pathogen recognition, anti-tumor immunity, responses to chromosomal double-stranded breaks (DSB), micronuclei surveillance, mitochondrial dysfunction, and autoimmunity (*4–10*). cGAS recognizes double-stranded DNA (dsDNA) in a sequence-independent, but length-dependent manner (*11, 12*). This allows cGAS to detect a broad spectrum of intracellular DNA anomalies while still maintaining regulatory control. That is, longer dsDNA fragments (≥45 base-pairs, bp), often indicative of pathogen invasion or genome instability, facilitate cGAS dimerization (*13–15*) and subsequent higher-order oligomerization into liquid-liquid phase separated (LLPS) condensates (*16–19*). Activated cGAS then synthesizes the secondary messenger, cyclic GMP-AMP (cGAMP), which binds to stimulator of interferon gene (STING) and triggers TANK-binding kinase (TBK1)-interferon regulatory factor 3 (IRF3) signaling to induce type I interferon (IFN-I) responses (*20–23*). Nevertheless, immunostimulatory DNA is rarely a perfectly contiguous duplex. Instead, it frequently harbors additional structural features, including supercoils, single-strand loops and gaps, unpaired Y-shaped ends and bubble-like regions (*24–30*). A previous study revealed that HIV cDNA-derived Y-form DNA activates cGAS, suggesting a role for such structural elements in immune signaling (*29*). Yet, the potential unique contributions of other structural motifs in the context of otherwise duplex DNA for cGAS activation are unknown.

Here, we identify dsDNA containing unpaired “bubble” regions as a potent stimulator for the cGAS-STING pathway. DNA bubbles are ubiquitously formed during the transcription and replication of both host and pathogenic DNA (*24, 31–33*). Enhanced cGAS activation is accompanied by stronger binding of cGAS to Bu-DNA than its fully complementary counterpart (C-dsDNA) and is independent of DNA topology and endogenous (such as mitochondrial DNA, mtDNA) or exogenous source. In a structure that deviates from canonical cGAS activation which involves dimers assembly along two long dsDNA fragment to form ladder-like higher-order oligomers (*15*) leading to condensate formation (*16–19*), our cryo-EM structure at 2.75 Å resolution, combined with biochemical assays and single molecule (sm) FRET experiments reveals a distinct mode of cGAS bending of Bu-DNA into a V-shape, using its unpaired region as a flexible hinge to yield a noncanonical 2:1 stoichiometry *in cis*.

### Bu-DNA is a more potent activator of the cGAS-STING pathway than fully duplex dsDNA

DNA bubbles, arising from dynamic and localized helix openings or “breathing”, are a ubiquitous structural feature of duplex DNA present in both endogenous and exogenous dsDNA (*24, 31–33*). These strand-separated regions represent common, but under-explored DNA structural motifs, whose role in innate immune signaling remains unclear. To address this directly, we synthesized an 88-bp duplex DNA containing a defined bubble region by introducing a 10-nucleotide (nt) mismatch (Fig. 1A). The presence of the unpaired region was validated using S1 nuclease digestion, which selectively cleaves single-stranded DNA. As expected, S1 nuclease fragmented Bu-DNA, confirming the accessibility of the bubble region, whereas a control, fully complementary dsDNA (C-dsDNA) remained intact (Fig. 1B).

**Fig. 1.**
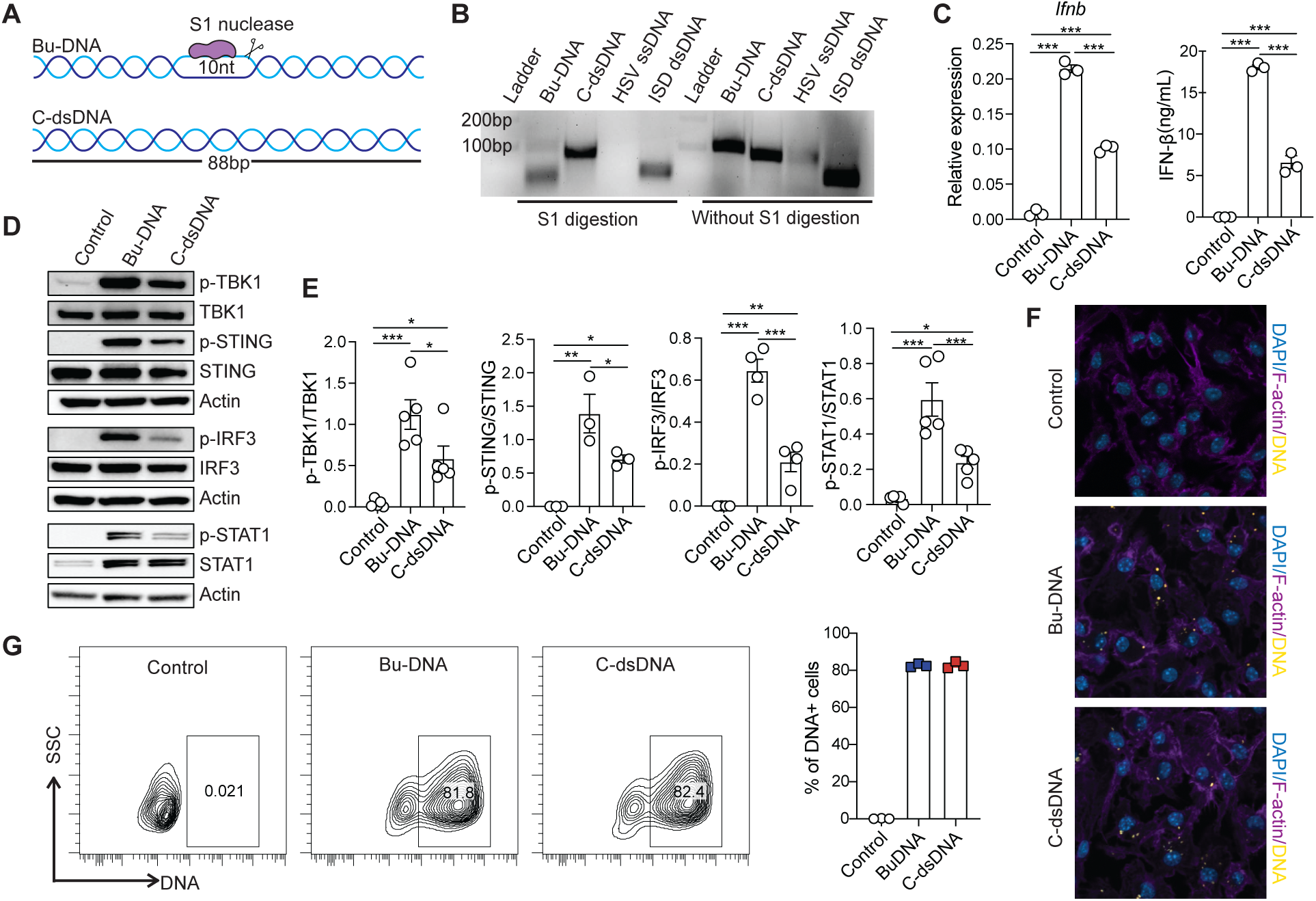
Bu-DNA enhances IFN-I responses. (**A-B**) Schematic of the 88-bp Bu-DNA and C-dsDNA (A) and representative gel validation of the structures following S1 nuclease digestion (B). At least three independent experiments were performed. HSV: herpes simplex virus, 100-nt ssDNA; ISD: interferon stimulatory DNA, 45-bp dsDNA. (**C-E**) mRNA (C, left) and protein (C, right) of IFN-β, and western blot analyses [(D), and densitometry in (E)] of WT BMDMs transfected with Bu-DNA, C-dsDNA for 6 or 18 hours, or left untreated (control). At least three independent experiments were performed with representative results shown in (C) and (D). Each dot in (E) represents one biological replicate in one independent experiment. Error bars depict the mean ± SEM. *p<0.05; **p<0.01; ***p<0.001. (**F**) Confocal microscopy of BMDMs transfected with Tetramethylrhodamine (TAMRA) labeled Bu-DNA, C-dsDNA for 6 hours, or left untreated (control), followed with DAPI (blue, for nuclear DNA) and phalloidin-AF647 (purple, for filamentous-actin) staining. (**G**) Flow cytometry analysis of BMDMs transfected with TAMRA labeled Bu-DNA, C-dsDNA for 6 hours, or left untreated as the control.

To evaluate the relative impact of Bu-DNA on innate immune signaling, we transfected equal amounts of Bu-DNA and C-dsDNA into bone marrow-derived macrophages (BMDMs) from wild-type (WT) mice. Bu-DNA triggered significantly more IFN-β production than C-dsDNA at both the mRNA and protein level (Fig. 1C) and further enhanced STAT1 phosphorylation (Fig. 1D, quantitated in Fig. 1E). Markedly stronger phosphorylation of TBK1, STING, and IRF3 (key downstream effectors of the cGAS-STING pathway) was also induced by Bu-DNA transfection compared to C-dsDNA (Fig. 1D, quantitated in Fig. 1E). Furthermore, transfection of Bu-DNA into human primary peripheral blood mononuclear cells (PBMCs) similarly resulted in elevated production of IFN-β (fig. S1A), indicating a conserved enhancement in both mouse and human cells. Consistent with this finding, higher IFN-β was observed in the human endothelial cell line EA.hy926 (fig. S1B), which is known to activate type I IFN signaling upon KSHV (Kaposi’s sarcoma-associated herpesvirus) virus infection (*34*). In addition, we also found an increase in pro-inflammatory cytokine IL-6 (fig. S1C) and elevated phospho-IκB upon Bu-DNA stimulation compared to C-dsDNA, although we did not observe a clear difference in phosphorylated p65 (*35, 36*) (fig. S1D, quantitation of multiple experiments in fig. S1E). The above differences were not attributed to transfection efficiency, as both confocal microscopy and flow cytometry demonstrated comparable cytosolic delivery of fluorescent-labeled Bu-DNA and C-dsDNA (Fig. 1, F and G).

To determine whether the enhanced IFN-I response to Bu-DNA was dependent on the cGAS-STING pathway, we assessed induction of cytokines in primary BMDMs derived from mice deficient in innate DNA sensing. Transfection of either Bu-DNA or C-dsDNA into cGAS-deficient (*Cgas^−/-^)* BMDMs failed to induce IFN-β although Bu-DNA induced more *Ifnb* and IFN-β than C-dsDNA in WT controls (Fig. 2A). Phosphorylation of TBK1, STING, IRF3 and STAT1 was also abolished in the absence of cGAS (Fig. 2B, quantitated in Fig. 2C), indicating that cGAS is essential for sensing cytosolic DNAs. Reconstitution of full-length mouse cGAS (cGAS^FL^) in *Cgas^−/-^* immortalized BMDMs (iBMDMs) restored the IFN-I responses, recapitulating the differential responses to Bu-DNA vs. C-dsDNA (Fig. 2D). Similarly, BMDMs lacking STING (encoded by *Tmem173*) were unable to express *Ifnb* in response to either form of DNA (Fig. 2E) and failed to show phosphorylation of STING-TBK1-IRF3 pathway components (Fig. 2F, quantitated in Fig. 2G), underscoring that both cGAS and STING are indispensable for sensing and responding to Bu-DNA to induce IFN-I responses. Similarly, the modest increase in IL-6 production upon Bu-DNA transfection was dependent on both cGAS and STING (*35, 36*) (fig. S1, F to L). Given that Toll-like receptors (TLRs) also mediate inflammatory responses via endosomal DNA sensing (*37–39*), we then examined *Myd88*^−/-^ BMDMs lacking the central TLR adaptor MyD88 (*37–39*). Here, Bu-DNA still triggered enhanced cytokine induction, like in WT cells, strongly suggesting that TLRs are not involved (Fig. 2H, and fig. S1M). It is noteworthy that cGAS can be found in the nucleus and remains inactive for cytokine induction, as its DNA-binding sites are shielded by histones (*40–46*). In these experiments, neither Bu-DNA nor C-dsDNA transfection penetrated the nucleus as revealed by DAPI co-staining, indicating that the observed inflammatory responses occurred in the cytosol (Fig. 1F). Based in these results, we conclude that Bu-DNA activates cGAS-STING signaling to a greater extent than fully duplex dsDNA.

**Fig. 2.**
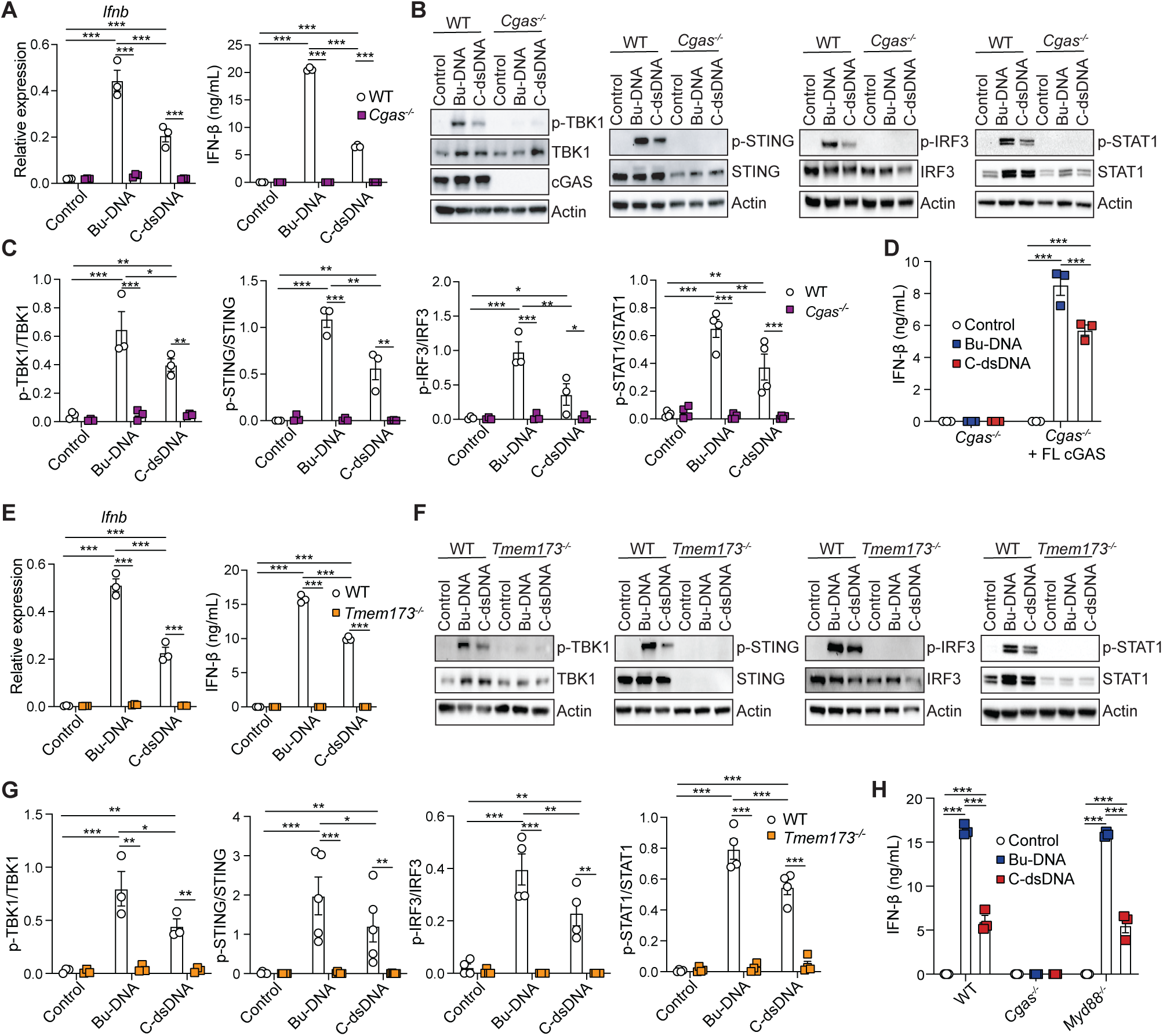
Bu-DNA-induced IFN-I responses are cGAS- and STING-dependent. (**A-C**) mRNA (A, left) and protein (A, right) of IFN-β, and western blot analyses [(B), and densitometry in (C)] of WT and *Cgas^−/-^* BMDMs transfected with Bu-DNA, C-dsDNA for 6 or 18 hours, or left untreated. (**D**) IFN-β in *Cgas^−/-^* iBMDM, unmodified or reconstituted with cGAS^FL^, transfected with Bu-DNA, C-dsDNA for 18 hours, or left untreated. (**E-G**) mRNA (E, left) and protein levels (E, right) of IFN-β, and western blot analyses [(F), and densitometry in (G)] of WT and *Tmem173^−/-^*BMDM transfected with Bu-DNA, C-dsDNA for 6 or 18 hours, or left untreated. (**H**) IFN-β in WT, *Cgas^−/-^*or *Myd88^−/-^* BMDMs transfected with Bu-DNA, C-dsDNA for 18 hours, or left untreated. Each dot in (C) and (G) represents one biological replicate in one independent experiment. At least three independent experiments were performed with representative results shown. Error bars depict the mean ± SEM. Comparisons are made within the same genotype of cells for (D) and (H). *p<0.05; **p<0.01; ***p<0.001.

### The presence of a bubble enhances cGAS sensing across different dsDNA topologies

Pathogenic dsDNA assumes various topologies in addition to simple linear duplexes. For instance, the genomes of papillomavirus and hepatitis B virus are circular (*47, 48*), and bacterial plasmids frequently exist in a supercoiled (SC) form (*49, 50*). To test whether the enhanced sensing of Bu-DNA by cGAS is influenced by overall dsDNA topology, we engineered two circular DNA constructs: the circular bubble DNA (cirBu-DNA) containing a 20-nt bubble and circular complementary DNA (cirC-dsDNA) (Fig. 3A). Transfection of cirBu-DNA into BMDMs elicited significantly higher production of IFN-β compared to cirC-dsDNA transfection (Fig. 3B). These responses were entirely dependent on both cGAS and STING as demonstrated in targeted gene-deficient cells (Fig. 3, C and D).

**Fig. 3.**
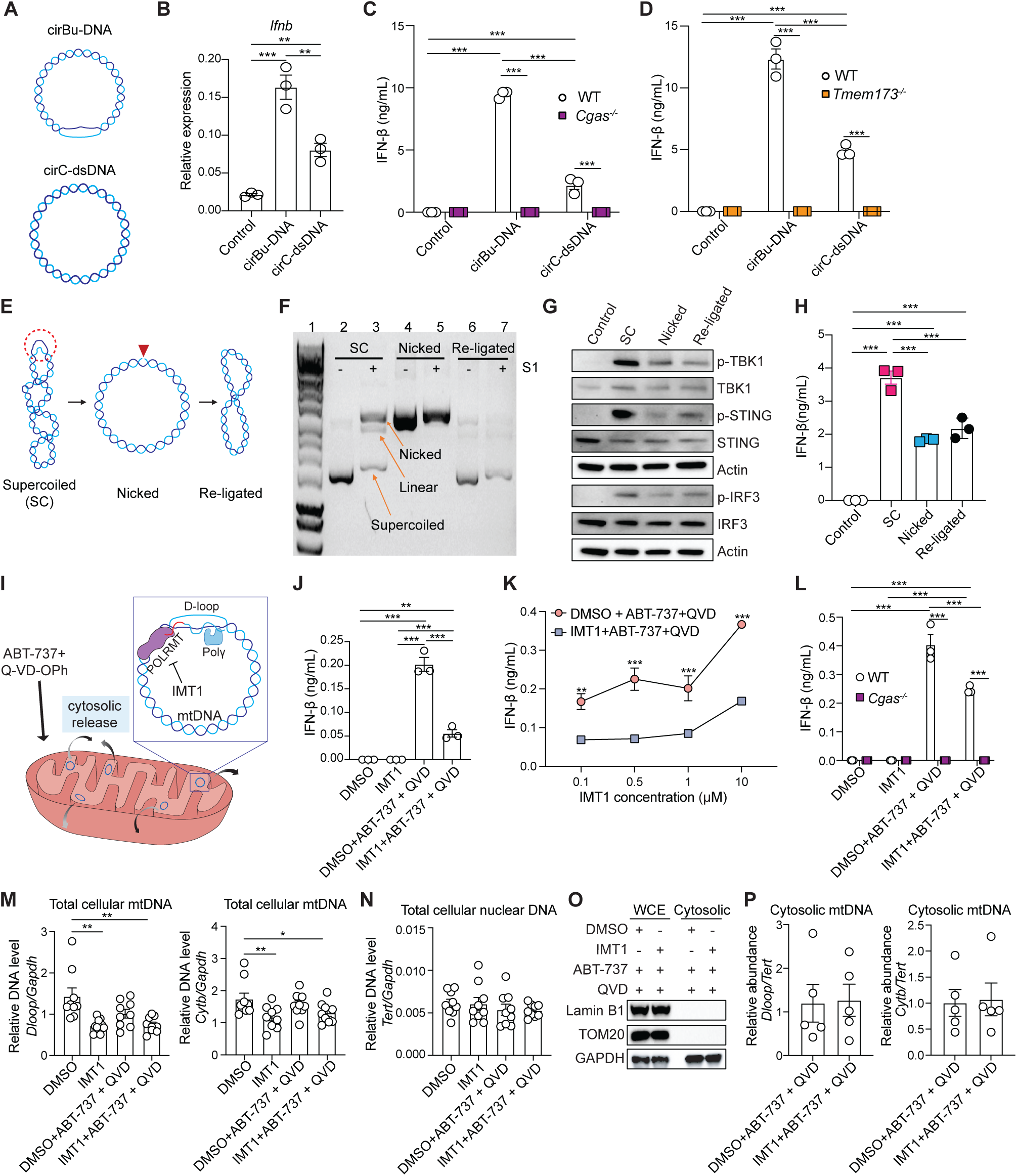
Bubble in circular DNA, supercoiled plasmid or mtDNA enhances cGAS-STING signaling. (**A-D**) Schematic of cirBu-DNA and cirC-dsDNA (A). mRNA (B) and protein levels (C-D) of IFN-β in WT, *Cgas^−/-^* (C) or *Tmem173^−/-^*(D) BMDMs transfected with the indicated constructs or left untreated. (**E**) Schematic of plasmid in supercoiled (SC), nicked (Nicked), and re-ligated (Re-ligated) forms. Red circle, unwinding region in SC. A nick was introduced (red inverted triangle) to SC and then re-ligated. (**F**) Gel electrophoresis of SC (lanes 2-3), nicked (lanes 4-5), and re-ligated (lanes 6-7) plasmids, with (+) or without (-) S1 nuclease digestion. Arrows denote SC, linear, and nicked plasmids. (**G-H**) Immunoblot (G) and ELISA (H) of WT BMDMs transfected with constructs in (E). (I) Experimental design: ABT-737 and Q-VD-OPh promote mtDNA release while IMT1 inhibits POLRMT and RNA primer synthesis (red line) for replication, thereby reducing D-loop formation. (**J-L**) IFN-β levels in WT or *Cgas^−/-^* iBMDMs upon IMT1 and ABT-737 + QVD treatment to cause cytosolic release of mtDNA. IMT1, 0.5 μM unless specified in (K). (**M-N**) Total cellular mtDNA and nuclear DNA with indicated treatments. (**O-P**) Cytosolic fraction verification and quantification of cytosolic mtDNA. Experiments were independently performed at least three times. Error bars depict the mean ± SEM. *p<0.05; **p<0.01; ***p<0.001.

While most DNA viruses replicate in the nucleus, vaccinia virus is an exception and replicates in the cytoplasm, where its DNA can activate cytoplasmic sensors (*51–53*). However, wild-type vaccinia virus encodes cGAS-STING inhibitors (*54, 55*), while modified mutant strains missing these are usually proliferation defective (*56, 57*). Thus, we searched for a different physiological model to circumvent these constraints. Plasmids are used clinically in DNA vaccines and gene therapies (*58, 59*). Supercoiled plasmids are susceptible to digestion by single-strand-specific nucleases (*60–64*) due to transiently unwound regions generated by torsional stress (*65–68*). As expected, treatment of pUC19 plasmids with S1 nuclease converted supercoiled DNA into relaxed/nicked circular and linear forms (Fig. 3F, lane 2 vs. 3). To prepare proper controls for testing these naturally occurring “bubbles”, we first introduced a single nick to generate relaxed circular DNA (Nicked, Fig. 3E), which alleviated torsional stress and rendered the plasmid largely resistant to further S1 digestion (Fig. 3F, lane 4 vs. 5). To further distinguish the effect of bubble from the nick, we re-ligated the nicked DNA using T4 DNA ligase (Re-ligated), restoring a closed circular DNA lacking torsional stress (Fig. 3E). The religated DNA was similarly resistant to S1 nuclease (Fig. 3F, lane 6 vs. 7). We then compared the immunostimulatory responses of these plasmid forms. Upon transfection, supercoiled plasmids induced the strongest cGAS-STING signaling (Fig. 3G) and triggered the highest levels of IFN-β compared to nicked or re-ligated plasmids (Fig. 3H).

Next, we asked whether endogenously generated Bu-DNA influences innate immune sensing. For this, we turned to mtDNA, which is a 16.5-kb circular molecule and known ligand for cGAS-STING and other innate immune inflammatory pathways (*1, 69, 70*). Release of mtDNA into the cytosol during certain viral infections and many other cellular stress conditions triggers cGAS-STING pathway and IFN-I responses (*69, 71–73*). The displacement-loop (D-loop) region of mtDNA is a stable, three-stranded structure formed during DNA replication, when the two strands of mtDNA are unwound and held apart by the nascent third strand of mtDNA that remains bound to the template strand. This results in a bubble-like configuration comprising a duplex (template and nascent strand) with a displaced single strand (non-template strand) (*69, 74*). We investigated whether the presence of the D-loop structure in mtDNA can enhance cGAS sensing. To suppress D-loop formation, we used IMT1, a selective small molecule inhibitor of the mitochondrial RNA polymerase (POLRMT) that does not affect cellular RNA polymerases (*75–77*). Since the RNA primers needed to initiate mtDNA replication are generated by POLRMT-mediated transcription (*69, 70, 78*), IMT1 will inhibit downstream D-loop formation (as illustrated in Fig. 3I). To promote mtDNA release into the cytosol and thereby activate cGAS, we treated iBMDM with the Bcl-2 family inhibitor ABT-737 together with the pan-caspase inhibitor Q-VD-OPh, which elicited IFN-β production (Fig. 3J) as described previously (*79–81*). Consistent with D-loop inhibition, markedly reduced IFN-β production was observed across all IMT1 concentrations tested (Fig. 3K; 0.5 μM IMT1 for the rest) compared to untreated controls and this was cGAS-dependent (Fig. 3L). To interpret these experiments, we assessed mtDNA abundance and release under these treatments. As expected that POLRMT inhibitor would impair mtDNA replication (*69, 70, 78*), IMT1 decreased total mtDNA content as measured by *Dloop* or *Cytb* levels (Fig. 3M), and the inclusion of the cytosolic release agents of ABT-737 and Q-VD-OPh showed a similar reduction trend (Fig. 3M). The nuclear DNA remained unaffected by IMT1 treatment (*Tert* level, Fig. 3N). We next verified the integrity of cytosolic fractionation (Fig. 3O) and determined the cytosolic mtDNA. Notably, the abundance of mtDNA released into the cytoplasm upon ABT-737 and Q-VD-OPh treatment was indistinguishable in the absence or presence of IMT1 (Fig. 3P). These results suggest that a structural configuration within mtDNA associated with the bubble-like, D-loop structure is particularly immunostimulatory, while inhibition of this structure reduced IFN-β induction by cGAS. Taken together, these findings demonstrate that the presence of unwound regions in dsDNA, whether synthetically introduced (linear Bu-DNA or cirBu-DNA) or naturally generated (SC plasmid DNA or mtDNA), potentiates cGAS-STING pathway activation.

### Bu-DNA binds to cGAS more tightly than C-dsDNA and suppresses condensate formation

To elucidate mechanisms underlying these observations in cells, we monitored the binding of recombinant mouse full-length cGAS^FL^ toward fluoresceine-amidite (FAM)-labeled Bu-DNA and C-dsDNA. Here, cGAS^FL^ bound Bu-DNA at least 2.5-fold more tightly in both direct and competition-based binding assays (Fig. 4, A and B, fig. S2A). Moreover, as expected from the higher affinity, Bu-DNA induced cGAS catalytic function as represented by nucleotidyl transferase (NTase) activity (*82*) more readily than C-dsDNA at sub-saturating concentrations (Fig. 4C). Consistent with our observations using recombinant proteins, biotinylated (Bio)-Bu-DNA pulled down more endogenous cGAS from BMDM lysates than Bio-C-dsDNA, and inclusion of unlabeled Bu-DNA competitor disrupted the interaction between Bio-C-dsDNA and endogenous cGAS to a greater extent (Fig. 4D). cGAS is composed of an intrinsically disordered N-terminal region and a highly conserved C-terminal domain, with the latter containing the NTase catalytic core for catalysis and the Mab21 domain for DNA recognition (*12–14, 19, 83*). The catalytic domain of cGAS^CAT^ maintained the higher binding-affinity and enzymatic activity toward Bu-DNA, indicating that the enhancement arises without involving the N-domain (Fig. 4, E and F, and fig. S2B). Moreover, consistent with our in-solution measurements, pull-down assays with immobilized Bio-Bu-DNA captured more recombinant cGAS^CAT^ protein than Bio-C-dsDNA (Fig. 4G). Unlabeled Bu-DNA also effectively displaced Bio-C-dsDNA bound to cGAS^CAT^, whereas unlabeled C-dsDNA failed to compete off Bio-Bu-DNA (Fig. 4G).

**Fig. 4.**
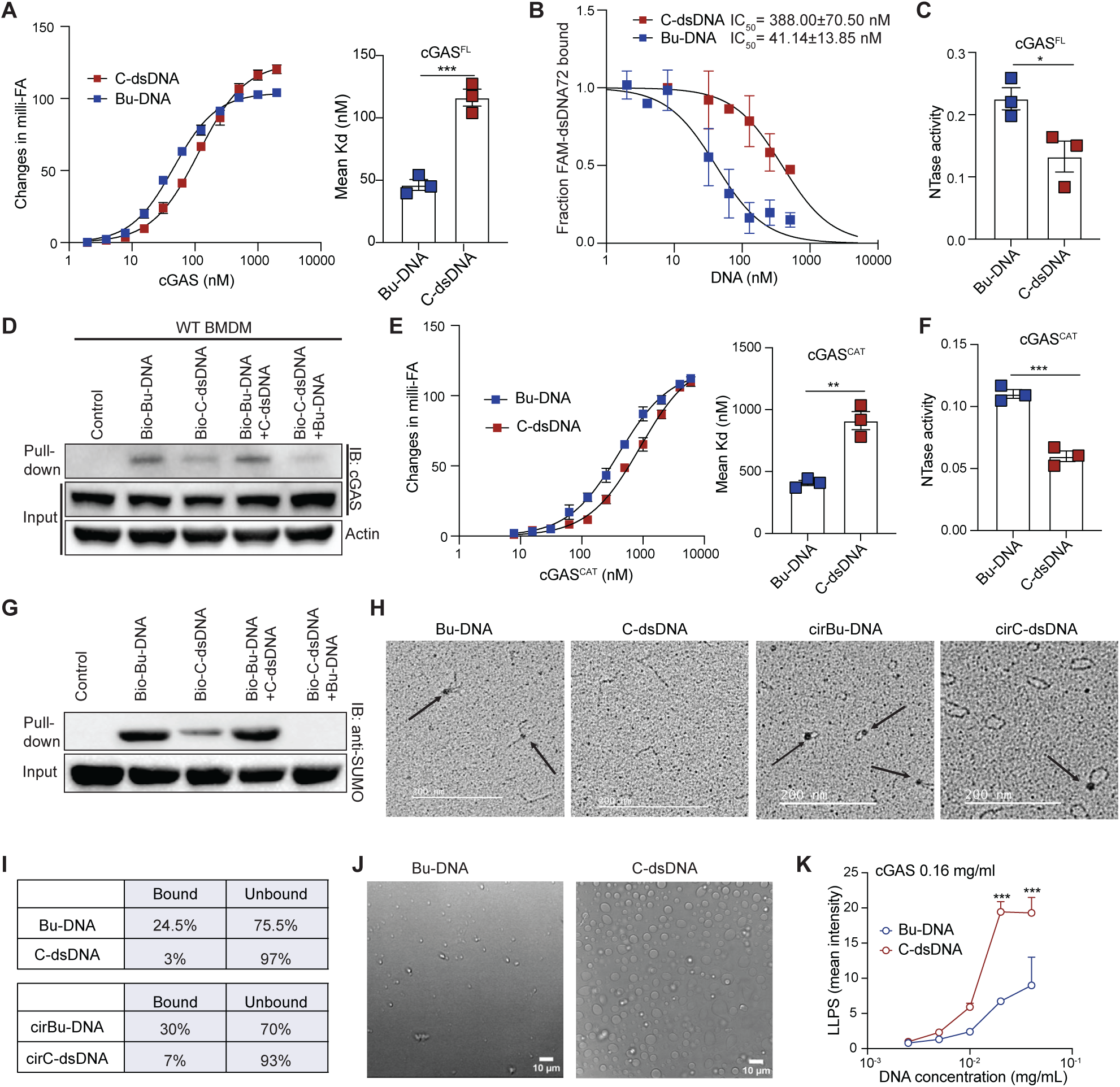
Bu-DNA exhibits higher binding affinity with cGAS than C-dsDNA. (**A**) FA of Bu-DNA vs. C-dsDNA binding to cGAS^FL^. Binding isotherm, left; Kd values, right. **(B)** Competition-based FA using 72 bp FAM-dsDNA and cGAS^FL^ with unlabeled Bu-DNA or C-dsDNA competitors. Fraction of bound FAM-dsDNA was plotted against competitor concentrations with IC_50_ values shown. **(C)** NTase activity of cGAS^FL^ with Bu-DNA or C-dsDNA. (**D**) Immunoprecipitation of endogenous cGAS from WT BMDM lysates by bio-DNAs with or without unlabeled competitors. (**E**) FA of Bu-DNA vs. C-dsDNA binding to cGAS^CAT^. Binding isotherm, left; Kd values, right. (**F**) NTase activity of cGAS^CAT^ with Bu-DNA or C-dsDNA. (**G)** Pull down of recombinant cGAS^CAT^ by bio-DNAs with or without unlabeled competitors. (**H-I**) EM images of cGAS^CAT^ binding to linear and circular DNAs (H) with arrows pointing to bound cGAS^CAT^ and percentages of bound vs. unbound events (I). **(J)** Bright-field view images of cGAS^FL^ incubated with Bu-DNA (left) or C-dsDNA (right). **(K**) Mean intensity of confocal images of cGAS^FL^-FITC with different concentrations of 60 bp HSV-derived Bu-DNA or C-dsDNA. Experiments were independently performed at least three times. Error bars depict the mean ± SEM (A, C, F and K); mean ± SD (B and E). *p<0.05, **p<0.01, ***p<0.001.

We next used transmission electron microscopy (TEM) to visualize complexes formed between the purified cGAS^CAT^ and either a 288 bp Bu-DNA containing a central 21 nt unpaired region, or its fully complementary counterpart (C-dsDNA) (Fig. 4, H and I). Longer DNA fragments were necessary for TEM visualization, and the use of cGAS^CAT^ minimized condensate formation that could otherwise obscure structural interpretation. Quantitative analysis revealed that cGAS^CAT^ bound Bu-DNA with markedly higher frequency (24.5%) compared to C-dsDNA (3%) (Fig. 4H, quantitated in Fig. 4I). A similar preference was observed using circularized DNA, with cGAS^CAT^ bound to 30% cirBu-DNA, but only 7% cirC-dsDNA (Fig. 4H, quantitated in Fig. 4I). Notably, cGAS^CAT^ particles (visualized as dark dots at arrows) were predominantly localized near the center of 288 bp Bu-DNA, corresponding to the bubble region (Fig. 4H). Taken together, these observations indicate that cGAS preferentially binds Bu-DNA compared to C-dsDNA.

Liquid-liquid phase separation (LLPS) is a critical mechanism underlying the activation and regulation of the cGAS-STING pathway (*16–19*). Thus, we next investigated whether the tighter binding of cGAS to Bu-DNA also promoted condensate formation, thereby facilitating its activation. Surprisingly, brightfield microscopy revealed that C-dsDNA formed far more condensates with cGAS^FL^ than Bu-dsDNA (Fig. 4J). To further characterize this behavior, we constructed a phase diagram and examined condensate formation across varying amounts of FITC-labeled cGAS^FL^ and DNAs (Fig. 4K, and fig. S2, C and D). Consistently, C-dsDNA induced more condensate formation than Bu-DNA (Fig. 4K, and fig. S2, C and D). Together, our results suggest that the enhancement of cGAS activation by Bu-DNA occurs through a mechanism that does not involve the promotion of condensate formation. In fact, Bu-DNA reduces condensate formation.

### cGAS bends Bu-DNA and forms a 2:1 dimer

To further elucidate how Bu-DNA preferentially binds and activates cGAS without promoting condensation, we determined the cryo-EM structure of the binary complex. We employed maltose-binding-protein (MBP)-tagged mouse cGAS^FL^, as we reasoned that the increased size might enhance the contrast for particle picking. Size exclusion chromatography of a 1:3 mixture of Bu-DNA with cGAS^FL^ yielded a single major peak with strong absorbance at 260nm (fig. S3B), indicating a stably bound complex. This peak fraction was used for cryo-EM grid preparation and data collection. 2D classification revealed that the vast majority of particles (91%) represent a single cGAS dimer (fig. S3C), whereas a much smaller fraction (9%) appeared to be the ladder-like, dimer-of-dimer arrangement (fig. S3C) as seen from the crystal structure of 39 bp dsDNA-bound cGAS^CAT^ (*15*) (PDB 5N6I). Given that the propagation of the ladder-like assembly has been implicated in driving the LLPS of cGAS•C-dsDNA complexes (*16, 19, 84*), our observations suggest that Bu-DNA enhances cGAS activity by largely constraining its oligomeric state to dimers without invoking condensation.

We refined the cryo-EM map corresponding to the predominant cGAS dimer species to an overall resolution of 2.75 Å. Here, the resulting model confirmed that the particles indeed corresponded to cGAS^CAT^ dimers assembled on Bu-DNA (Fig. 5, A and B, fig. S3, A and E, and movie S1). Notably, the N-domain of cGAS and the MBP tag were not resolved in the density map, suggesting that these regions are flexibly tethered (Fig. 5A). These observations are also consistent with our biochemical data indicating that cGAS^CAT^ itself maintained the higher binding-affinity toward Bu-DNA (Fig. 4, and fig. S2B). The interactions between cGAS side-chains and the contiguous duplex part of Bu-DNA are essentially the same as the previously determined crystal structures of cGAS^CAT^ dimers formed over two separate dsDNA fragments (2:2 dimers (*13–15*); fig. S4A). Notably, instead of the canonical 2:2 arrangement, cGAS dimerized on a single Bu-DNA fragment by sharply bending it into a V-shape (∼140°), using the central unpaired region as a flexible hinge (Fig. 5C).

**Fig. 5.**
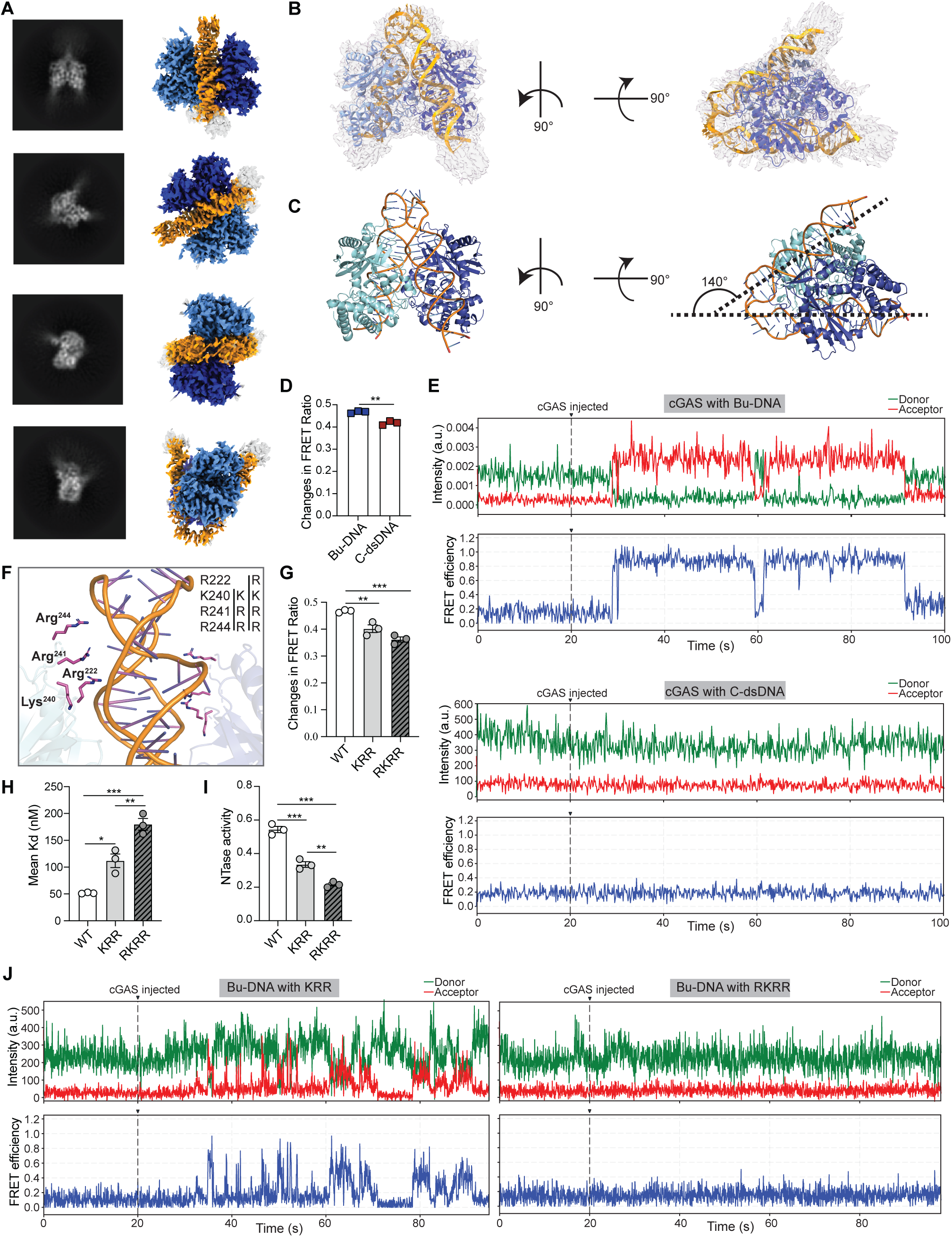
cGAS bends Bu-DNA into a V-shape with hinge-interacting residues crucial for the bending, binding and activity. (**A)** Representative 2D class averages (left) and corresponding 3D reconstruction (right) of cGAS-Bu-DNA complex. (**B-C**) Atomic model of V-shaped cGAS-Bu-DNA complex fitted into the 3D cryo-EM density map (B) or highlighted for Bu-DNA bent at an angle of ∼140° (C). Dark and light blue, two cGAS monomers; Bu-DNA, yellow; light grey, fitted model. (**D)** Changes in FRET ratio of Bu-DNA vs. C-dsDNA binding to cGAS^FL^ (see fig. S5A for the assay illustration). (**E**) Representative smFRET trace showing donor (green) and acceptor (red) fluorescence intensities and calculated FRET efficiency (blue) of cGAS with Bu-DNA (top two panels) or C-dsDNA (bottom two panels). FRET efficiency were calculated as intensity of (acceptor/(donor + acceptor) (see fig. S5B for the assay illustration). (**F**) Close-up view of cGAS and Bu-DNA interaction, with cGAS residues and Bu-DNA hinge regions highlighted. (**G-I**) Changes in the FRET ratio (G), FA assay (H), and NTase activity (I) of Bu-DNA binding to WT cGAS versus two cGAS mutants. (**J**) smFRET of two cGAS mutants with Bu-DNA.Experiments were independently performed at least three times (D-J). Error bars depict the mean ± SEM. *p<0.05; **p<0.01; ***p<0.001.

The observed dimer of dimers in our cryo-EM samples suggest that this subpopulation is likely responsible for the residual condensate forming activity of Bu-DNA. To determine whether the bubble region is involved in forming the canonical arrangement, we determined the cryo-EM structure corresponding to these particles (3.5Å resolution; Fig. S3F and fig. S4B). Our resulting model showed that two cGAS dimers are indeed bound to two contiguous dsDNA fragments in the same ladder-like manner seen from the previously reported crystal structure (fig. S4, B and C). Notably, the two dimers are formed exclusively over the 45-bp dsDNA arm of Bu-DNA without involving the central unpaired bubble region and the shorter arm (33-bp; fig. S3, C and D and fig. S4B). Also of note, as seen from the particles showing cGAS dimers, the cGAS N-terminal domain and the MBP tag along with the remaining portions of Bu-DNA remain unresolved, indicating that cGAS^CAT^ dominates the stable interactions with dsDNA in all cases. Together, our results revealed that Bu-DNA promotes *in cis* formation of active cGAS dimers on a single DNA fragment, obviating the canonical requirement for *in trans* assembly on two separate dsDNA fragments.

Our cryo-EM structure suggests that cGAS engages Bu-DNA by exploiting the flexibility of the unpaired region, which acts as a hinge to facilitate or stabilize a sharply bent, V-shaped DNA conformation. To further test this mechanism, we performed both bulk-solution and single-molecule Forster Resonance Energy Transfer (FRET) measurements using donor- and acceptor-labeled Bu-DNA_(D/A)_ and C-dsDNA_(D/A)_, (fig. S5, A and B). In bulk assays, Bu-DNA_(D/A)_ showed significantly higher FRET efficiency than C-dsDNA_(D/A)_ upon introducing cGAS^FL^ (Fig. 5D). In single-molecule (sm) experiments, a rapid transition from low to high FRET was observed from immobilized Bu-DNA_(D/A)_ upon introducing cGAS, with clear anti-correlated changes in donor and acceptor intensities (Fig. 5E). smFRET changes for multiple traces directly correlated with cGAS injection, as visualized in rastergrams (Fig. 5E and fig. S5C) and the raw movie (movie S2), indicating that cGAS consistently bends Bu-DNA. By contrast, C-dsDNA_(D/A)_ remained in the low-FRET state upon adding cGAS (Fig. 5E and fig. S5D, see movie S3). These results strongly suggest that cGAS directly bends Bu-DNA.

While inspecting our cryo-EM structure, we noted that the bent architecture appeared to be stabilized by several positively charged residues positioned at the edge of cGAS^CAT^ (Fig. 5F), which are often poorly resolved in previously determined crystal structures as employed dsDNA fragments are too short to reach these distal regions (*14*) (18 bp, PDB 4LEZ). To test whether these cGAS residues are essential for bending Bu-DNA, we generated two sets of charge-reversal mutations on cGAS^FL^ (positive to negative; Fig. 5F). Compared to WT, both mutants showed reduced bulk FRET efficiency (Fig. 5G), and concomitantly impaired binding and catalytic activities induced by Bu-DNA (Fig. 5, H and I). Moreover, in smFRET experiments, the KRR mutant showed far fewer traces indicative of bending events, and the RKRR mutant failed to bend Bu-DNA_(D/A)_ altogether (lack of smFRET signals, Fig. 5J and fig. S5E, see movie S4 and movie S5). These results corroborate our cryo-EM structure and demonstrate that cGAS can exploit the flexibility of DNA bubbles to promote the formation of active dimers.

## Discussion

The presence of dsDNA in the cytosol signals various intracellular crises, such as pathogen invasion, damaged or malfunctioning organelles (e.g. nucleus, mitochondria, or lysosomes), genotoxic stress, uncontrolled cell division, and engulfment of necrotic cells (*1–3*). Mislocalized, rogue dsDNA is recognized by cGAS and initiates STING-dependent activation of IFN-I responses (*20–23*). In addition to simple linear dsDNA, DNA frequently assumes various additional conformations and local structures, including circular, supercoiled, gapped, and bubbled forms (*24, 26, 30*). Currently, the mechanisms by which cGAS interacts with these nonlinear or non-contiguous dsDNAs remain under-appreciated and under-explored. Here, we reveal that dsDNA containing bubbles is a novel DNA form that hyperactivates cGAS-STING signaling. cGAS preferentially recognizes Bu-DNA (Fig. 4) and this interaction lowers the threshold for initiating cGAS-STING signaling, thereby eliciting stronger inflammatory responses than C-dsDNA (Fig. 1 and 2 and fig. S1). A similar enhancement was observed with human cGAS, as both primary human PBMCs and a human endothelial cell line produced higher level of IFN-β in response to Bu-DNA transfection (fig. S1, A and B). Mechanistically, although the canonical activation of cGAS entails 2cGAS:2dsDNA dimerization and subsequent condensation with contiguous dsDNA (*13–15*), we find that cGAS sharply bends Bu-DNA into a V shape to form a hyper-stable 2cGAS:1dsDNA dimer without invoking condensate formation (Fig. 4, Fig. 5 and fig. S2).

DNA bubbles are ubiquitous in biology, as transient unwinding of the double helix occurs spontaneously (*24, 31, 32*). Bacterial plasmids, for instance, are predominantly supercoiled and contain bubbles due to torsional stress (*49, 50*). Replication bubbles also occur during mtDNA (*70, 74, 85*) and viral DNA replication (e.g., HSV, HPV and HBV)(*47, 48, 86*). Furthermore, genome instability and defective mismatch repair can result in the accumulation of cytosolic dsDNA (*1, 7, 84, 87–89*) that may contain unpaired regions. Our findings demonstrate that cGAS preferentially respond to bubble-containing regions regardless of the overall dsDNA topology (Fig. 3). Compared to linear dsDNA forms, those that contain bubbles in circle or supercoiled plasmids enhance cGAS-STING and IFN-I responses. Interestingly, the stable D-loop form of mtDNA that is generated during mtDNA replication that has been appreciated for decades (*70*) is an endogenous bubble-like structure that also contributes to enhanced cGAS sensing when released into the cytoplasm (Fig. 3). Consistent with this, based on qPCR of purified cytosolic fractions, the mtDNA D-loop region is over-represented in the cytosol of *Tfam*-deficient cells that exhibit enhanced mtDNA release and cGAS-Sting activation (*71*). These results indicate that flexible regions within DNA may provide an attractive recognition motif within PAMPs and DAMPs for enhanced host innate immune responses. Conversely, since DNA bubbles also occur frequently in the host chromosomal DNA, how does the innate immune system prevent cGAS from reacting to these self-derived “benign” bubbles? One mechanism is the tethering of nuclear cGAS to nucleosomes, which restricts its activation (*40–46*). Importantly, the cGAS residues involved in bending Bu-DNA are also recognized to be involved in binding the acidic patch of nucleosomes (Fig. 5) (*40–45*). Thus, we surmise that the chromatin sequestration and tethering of cGAS effectively suppress spurious activation by Bu-DNA in the host nucleus.

Taken together, our study highlights the versatility of host defense strategies where cGAS can identify and adapt to different structural features as a pattern within PAMPs and DAMPs to tailor inflammatory responses. Our findings offer a new framework for deciphering how structural elements embedded in pathogenic- or damaged- nucleic acids can regulate PRR activity. Future studies will delineate other PRRs recognition of various DNA features at the molecular level and define how innate immunity interprets such cues to maintain a balance between defense and tolerance.

## Acknowledgement

Flow cytometry analysis was performed at the University of North Carolina at Chapel Hill (UNC-CH) flow cytometry facility. Confocal microscopy was conducted in the Department of Pathology and Laboratory Medicine at the UNC-CH. We thank Pablo Ariel for assistance in confocal microscopy and members of flow cytometry facility for help. Both UNC flow cytometry facility and MSL are supported in part by P30 CA016086 Cancer Center Support Grant to UNC Lineberger Comprehensive Cancer center. All cryo-EM data were collected at the Beckman Center for Cryo-EM at Johns Hopkins.

## Author contributions

S.Y. and S. Wu contributed equally to this work. J.S. and J.P.-Y.T. jointly supervised the project. S.Y., S. Wu, J.S. and J.P.- Y.T. designed and performed most experiments, analyzed data and wrote the manuscript with input from all authors. S.C. conducted phase separation studies, confocal microscopy and protein purification. X.L. made the initial discovery, generated cGAS constructs and purified cGAS-Cat protein. I.C. performed smFRET studies and analyzed the data. S. Willcox prepared circular DNAs and contributed to electron microscopy studies. K.C.B. generated wild-type and *Cgas^−/-^* iBMDM cells, made various cGAS constructs, established reconstituted iBMDM cell lines, and assisted with experimental design. X.H. performed and analyzed the EMSA assays. G.H. and W.J.B assisted with mouse breeding and reagent preparation. J.A.D. contributed to protein purification and experimental design. W.C. participated in experimental design and data analysis. P.L. provided *Cgas^−/-^* mice and helpful discussion. W.G.F provided assistance with protein purification. S.T. conducted the bright-field imaging of phase separation. G.S. supported the mtDNA studies and participated in experimental design and data analysis. G.D.B. supervised the smFRET studies. J.D.G. helped prepare circular DNAs, conducted electron microscopy, acquired representative images and contributed to experiment design.

## Data, materials and software availability

The model of the cGAS dimer-BuDNA88 complex has been deposited in the RCSB PDB (identifier 9DCK). The CryoEM density map has been deposited in the EMDB with the identifier EMDB-46754. The model for the cGAS tetramer-BuDNA88 complex has been deposited in the RCSB PDB (identifier 9OH4). The CryoEM density map has been deposited in the EMDB with the identifier EMDB-70483. All other EM data are available from the corresponding authors upon request.

## Supplementary Materials

Materials and Methods

Figures S1 to S5

Tables S1 to S2

Movies S1 to S5

