## Supplementary Materials for "cGAS bends unpaired DNA to form an unconventional structure that hyperactivates the innate immune response"

**The PDF file includes:**

Materials and Methods
Figs. S1 to S5
Tables S1 to S2

**Other Supplementary Materials for this manuscript include the following:**

Movies S1 to S5

### Materials and Methods

Mice, cell culture and transfection

Wild-type (C57BL/6), *Cgas*$^{-/-}$, *Tmem173*$^{-/-}$, and *Myd88*$^{-/-}$ mice were housed and bred under specific pathogen-free conditions at the animal facility of University of North Carolina at Chapel Hill, with protocols approved by the Institutional Animal Care and Use Committee (IACUC). Bone marrows were isolated from femurs and tibias of 6- to 10-week-old mice and differentiated into macrophages (BMDM) in DMEM complete medium with 10% FBS, L-glutamine, penicillin-streptomycin, non-essential aminol acid, sodium pyruvate and 30% L929 cell culture medium for 6-7 days. Wild-type and *Cgas*$^{-/-}$ immortalized bone marrow derived macrophages (iBMDMs) were generated from isolated bone marrow using J2 retrovirus from CREJ2 cell supernatants as previous reported *(90)*. Human peripheral blood mononuclear cells (hPBMC) were isolated from primary human leukopak (Gulf Coast Blood) using Ficoll®-Paque PREMIUM density gradient (Millipore Sigma). Following isolation, CD14$^{+}$ monocytes were negatively selected using Dynabeads$^{TM}$ Untouched$^{TM}$ Human Monocytes Kit (ThermoFisher 11350D) according to the manufacturer's protocol and cultured in RPMI complete medium supplemented with 10% FBS, L-glutamine, penicillin-streptomycin, non-essential aminol acid, sodium pyruvate. Human endothelial cell line EA.hy926 was a gift from Dr. Pengda Liu's lab and maintained in complete DMEM medium. Transfection of DNAs (2 µg/mL) was performed using Lipofectamine 3000 Transfection Reagent (ThermoFisher) following the manufacturer's protocol for 6 or 18 hours.

DNAs

88 bp DNAs: The forward and reverse sequences were synthesized by Integrated DNA technologies (IDT) and annealed at 95°C with 5°C decrease per minute until 4°C. Bu-DNA and C-dsDNA shared the same forward sequence while the reverse ones differed (table S1). 288 bp

DNAs and 170 bp circular DNAs were synthesized by IDT and used for electron microscopy (sequences in table S1). To make cirBu-DNA and cirC-dsDNA, equal amounts of forward and reverse 170nt oligos were mixed at 5 nM concentration in 30 mL buffer containing 10 mM Tris, 1 mM EDTA and 50 mM NaCl, which were distributed in small tubes up to 1.5 mL each, heated to 95°C for 5 min in a water bath and slow cooled in a bucket until reaching room temperature. The annealed mixture was pooled together and concentrated using Microsep Centrifugal Devices (10K, Pall Corporation) until volume within the 100-200 μL range. The annealing buffer was exchanged with water (2 x 1 mL) using the same centrifugal devices over two more centrifugations. The clean retentates were removed and placed in an eppendorf tube for ligation. T4 ligase buffer and immobilized T4 ligase beads were added to the DNA and incubated for 30 min at room temperature. The magnetic beads were then separated from the solution using a magnetic stand and the clear DNA solution was further cleaned on a Zymoclean column (DNA Clean & Concentrator, Zymo Research). DNA was eluted in water and the circularization efficiency was evaluated by EM.

S1 nuclease digestion, nick creation and ligation

To validate bubble formation, S1 nuclease (ThermoFisher Scientific) was employed at room temperature for 30 min and deactivated at 70°C for 10 min with addition of EDTA according to the manufacturer's protocol. The digested products were then examined by 1.5-3% agarose gel electrophoresis. Similarly, pUC19 plasmids were digested with S1 nuclease to examine the existence of bubbles in supercoiled plasmids. To create a nick in supercoiled DNA, pUC19 plasmids were digested by Nt.BspQI enzyme at 50°C for one hour and then cleaned up using DNA clean-up kit (DNA Clean & Concentrator, Zymo Research). Then, nicked plasmids were ligated using T4 ligase at RT for one hour. The reaction was deactivated at 65°C for 10min and cleaned up using DNA clean-up kit.

Flow cytometry

BMDMs transfected with or without Tetramethylrhodamine (TAMRA)-labeled DNAs were resuspended in FACS buffer and fixed with 4% paraformaldehyde for 15 min. After washing and centrifuging, samples were assayed on a ThermoFisher Attune NxT flow cytometry machine at UNC Flow Cytometry Core Facility. Acquired data was analyzed with FlowJo software and plotted in Adobe Illustrator 2025.

RNA extraction and real-time PCR

Cells were transfected with DNAs for 6 hours. Total RNA was extracted from cell lysates using the RNeasy Mini Kit (74106, Qiagen) according to the manufacturer's protocol. cDNA was synthesized using iScript™ cDNA synthesis kit (Bio-rad). Real-time PCR analysis was performed using the iTag Universal SYBR Green Supermix reagent (Bio-Rad) on a QuantStudio 6 Real Time PCR System (Applied Biosystems). Gene expression levels were normalized to *Actb* or *b2m* expression and analyzed using the $\Delta$ Ct method. Primer sequences are listed in table S1.

Enzyme-linked immunosorbent assay (ELISA)

Cytokines IL-6 in the supernatants of transfected cells (18 hours) were measured according to manufacturer's protocols (BioLegend). IFN-β was quantified using anti-mouse IFN-β capture antibody (Santa Cruz Biotechnology, sc-57201), detection antibody (PBL Assay Science, 32400), and mouse IFN-β protein standard (BioLegend, 581309).

Western blot analysis

Cells were transfected with DNAs for 6 or 18 hours and lysed in RIPA buffer supplemented with PhosStop phosphatase inhibitor (Roche) and complete EDTA-free protease inhibitor (Roche) and denatured in Laemmli sample buffer. Proteins were separated by SDS-PAGE in 4-12% NuPAGE Bis-Tris gel (Invitrogen) and transferred to Nitrocellulose membrane, 0.2μm (Bio-rad). Membranes were blocked with TBST buffer containing 5% nonfat milk and incubated with primary antibody overnight for specific targets, including phosphorylated (p-)STING (72971), STING (13647), p-TBK1 (5483), TBK1 (3504), p-IRF3 (4947), IRF3 (4902), cGAS (31659), p-p65 (3033), p65 (8242), p-STAT1 (9172), STAT1 (9167), p-IκB (2859), and IκB (4812) (Cell Signaling Technology), followed by HRP-conjugated secondary antibodies (Jackson ImmunoResearch). Cells were transfected with DNAs for 6 hours for the analysis of STING, TBK1, IRF3, and cGAS and for 18 hours for STAT1, p65, and IκB. Signals were detected by the ChemiDoc Imaging system (Bio-Rad) and quantified using ImageJ.

Cytosolic DNA isolation and mtDNA quantitation

To study mtDNA, WT and *Cgas*$^{-/-}$ iBMDM cells were treated with DMSO, 0.5μM IMT1 (MedChemExpress, HY-134539), 10 μM ABT-737 (MedChemExpress, HY-50908) or 10μM Q-VD-OPh (MedChemExpress, HY-12305) as indicated for 18 hours. DNAs were extracted and purified using DNeasy blood and tissue kit from Qiagen and then analyzed by real-time PCR for total cellular mtDNA (*Dloop*, *Cytb*) or nuclear DNA (*Actin*, *Tert*) normalized to *Gapdh* expression. The cytosolic DNA isolation was conducted as previous reports *(71, 81)*. Briefly, 12 x $10^6$ iBMDM were split into two parts, one as whole cell lysate (WCE) and the other as cytosolic part. For isolating DNA from WCE, cells were pelleted and lyzed with SDS lysis buffer (20mM Tris, pH8, 1% SDS, protease inhibitors) at 95°C for 15 mins. For Cytosolic fraction, membranes were selectively permeabilized with 18ug/mL digitonin (50mM HEPES, pH 7.4, 150mM NaCl, protease inhibitor) for 20 mins at 4°C. Supernatants were collected after spinning down to remove any pellets. 50 μL supernatant from WCE or cytosolic fraction were run for western blot analysis and the remaining 450ul for phenol-chloroform isolation of DNA. Isolated DNAs were analysis by real-time PCR for mtDNA presence (*Dloop, Cytb* relative to *Tert* expression) and normalized to WCE part.

cGAS-DNA EMSA assay

Bu-DNA or C-dsDNA (20ng) were mixed with recombinant $cGAS^{FL}$ (0, 0.0161 µg, 0.03125 µg, 0.0625 µg 0.125 µg, 0.25 µg) or $cGAS^{CAT}$ protein (0, 0.25 µg, 0.5 µg, 1 µg, 2 µg, 4 µg) in buffer containing 10 mM HEPES (pH 7.3), 20 mM KCl, 1mM MgCl2, 1mM DTT and incubated at room temperature for 30 min. EMSA loading buffer was then added to the reaction and immediately loaded to native acrylamide gel (Novex TBE gel, 4-12%, EC62352BOX, Invitrogen). After 40 min electrophoresis, gels were stained with SybrGold (S11494, Invitrogen) for 15 min and band intensities determined by imaging (Bio-Rad Laboratories). The free DNA intensity was quantified through ImageJ. The Kd was calculated from the intensity data using GraphPad Prism10.

Confocal microscopy

BMDMs were plated on coverslips and transfected with fluorescent-labeled Bu-DNA or C-dsDNA for 3 hours. After PBS washes, the coverslips were fixed with 2% paraformaldehyde for 20 mins at room temperature and permeabilized in blocking buffer (0.1% saponin, 50mM $NH_4Cl$, and 2% goat serum) for 30 mins in the dark, followed by DAPI and Phalloidin Alexa Fluor Plus 647 (Invitrogen, A30107) staining for one hour. Coverslips were mounted and visualized at the UNC-CH Microscopy Services Laboratory (MSL) using a Zeiss LSM900 confocal microscope with a 40x/1.4 Oil Plan Apo objective.

cGAS protein purification

$cGAS^{FL}$ and $cGAS^{CAT}$ were expressed and purified as previously described *(14, 18)*. Briefly, BL21 Star (DE3) chemically competent *E. coli* cells transformed with pET-28a-SUMO plasmid containing 6xHis-Sumo-cGAS-FL and 6xHis-Sumo-cGAS-CAT, were cultured in 2 L flasks with 1 L LB medium at 37°C shaking at 250 rpm until OD600 reached a value of 0.6. Then, the bacterial culture was cooled to 16°C and IPTG was added to the medium at a final concentration of 0.5 mM. After 16-20 hours, the culture medium was centrifuged at 4,000 x g at 4°C. The bacterial pellet was collected, resuspended in PBS buffer containing 300 mM NaCl, 1 mM PMFS, 0.1 mg/ml lysozyme, and 1 U/ml benzonase, and then sonicated. Cell lysate was centrifuged at 30,000 x g at 4°C. Supernatant was applied to the TALON metal affinity resin in gravity columns, which was washed with PBS buffer with 10 mM imidazole and incubated with PBS buffer containing ULP1 sumo protease at 4°C overnight. After cleavage, cGAS protein was washed off the resin and purified using a Heparin column with a linear NaCl gradient elution. The eluted protein was concentrated and applied to Superdex 200 size exclusion column in a buffer containing 20 mM HEPES (pH 7.5) and 150 mM NaCl. The final purified protein was concentrated and stored at -80°C freezer.

For cryo-EM, NTase assay, FRET assays and FA assays, proteins were purified as below. Full-length and catalytic domain (residue 147-507) of mouse cGAS were cloned into the pET28b vector (Novagen) with an N-terminal 6×His-MBP-tag containing a TEV protease cleavage site. Plasmids encoding wild-type and mutant cGAS constructs were expressed in E. coli BL21-(DE3) cells at 16°C using 0.3 mM IPTG for 18 hours. Cells were lysed by sonication in the lysis buffer (20 mM Na-phosphate at pH 7.5, 500 mM NaCl, 40 mM imidazole, and 0.5 mM TCEP) and purified by Ni-NTA chromatography. The fusion protein was cleaved by TEV protease incubation at 4°C overnight, and the MBP tag and uncleaved fusion protein were removed using the Ni-NTA and amylose columns. The untagged protein (flow-through) was further purified by size-exclusion chromatography (Superdex 200 for full-length, Superdex 75 for catalytic domain, Cytiva) and concentrated in buffer with 20 mM Tris-HCl (pH 7.5), 150 mM NaCl, 10% glycerol and 0.5 mM TCEP. All purified cGAS proteins were flash-frozen in liquid nitrogen and stored at -80°C.

Liquid-liquid phase separation (LLPS)

Broad field images of condensates were collected at 25 ± 2° C with 0.075mg/mL DNAs and 0.174 mg/mL cGAS-FL protein in the buffer containing 25 mM Tris-acetate (pH 7.4), 125 mM potassium acetate (pH 7.4), 1 mM TCEP, 5 mM Mg(acetate)2, and 5% glycerol. Confocal microscopy analysis of LLPS was conducted using FITC-labeled $cGAS^{FL}$. Briefly, DNAs were serially diluted and incubated with various concentrations of FITC-$cGAS^{FL}$ protein in a 384-well plate at 37°C using the protein purification buffer. After protein-DNA incubation, LLPS between FITC-$cGAS^{FL}$ with Bu-DNA or C-dsDNA were visualized using a confocal microscope (Zeiss LSM 900) with images taken at the UNC-CH Microscopy Services Laboratory (MSL).

Pulldown assay

Biotin-labeled Bu-DNA or C-dsDNA were first immobilized to streptavidin beads and then incubated with BMDMs lysed in NP-40 buffer (Boston Bioproducts) containing protease inhibitor cocktail for three hours. The beads were washed at least 5 times and eluted with 1x Laemmli sample buffer at 96°C for 5 min, followed by western blot analysis using anti-cGAS antibody (Cell signaling Technology). For the competition assay, purified mouse $cGAS^{CAT}$ with the SUMO tag was first incubated with or without unlabeled competitor DNAs for one hour before incubating with biotinylated Bu-DNA or C-dsDNA immobilized on streptavidin beads. Anti-SUMO antibody (Sigma) was applied for western blot analysis of $cGAS^{CAT}$.

Transmission electron microscopy (TEM)

DNA samples were mixed with a buffer containing spermidine and adsorbed onto carbon-covered copper grids glow-charged shortly before sample application. After adsorption of the samples for

2–3 min, the grids were washed with EM-grade water and dehydrated through a graded ethanol series from 25% to 95%. Following rapid air-drying, the grids were rotary shadow cast with tungsten at $2 \times 10^{-6}$ torr of pressure. Samples were examined by an FEI T12 TEM scope equipped with a Gatan 2kx2k Orius CCD camera at 40 kV.

Pyrophosphatase-coupled cGAS activity assay

The activity of cGAS (NTase) was measured using the pyrophosphatase-coupled NTase assay. Briefly, 100 nM mouse cGAS was incubated with 50 nM of *E. coli* pyrophosphatase, 200 μM of substrates ATP/GTP plus 88 bp Bu-DNA or C-dsDNA in the reaction buffer [(25 mM Tris-acetate (Ac) pH 7.4, 125 mM KAc, 1 mM TCEP, 5 mM $Mg(Ac)_2$, pH 7.4, and 5% glycerol)] at 25 ± 2°C (RT) for three hours. DNA concentration was at 6.25 nM (~ 0.07 ng/ μL) and 12.5 nM (~ 0.14 ng/ μL) (DNA concentrations were normalized based on cGAS binding site, assuming one binding site per 18 bp of DNA) for WT $cGAS^{FL}$ and $cGAS^{CAT}$, respectively. DNA concentration was fixed at 50 nM (~ 0.56 ng/ μL) for all the mutated $cGAS^{FL}$. 88 bp Bu-DNA and C-dsDNA were annealed using the oligo sequences described above. The reaction was quenched with an equal volume of the quench buffer (reaction buffer plus 25 mM EDTA) in a 384-well plate. Quenched reactions were mixed with 10 μL malachite green color development solution and developed over 45 min at RT. The absorbance at 620 nm was recorded using a Tecan M1000 plate reader. The NTase activity was normalized to control reactions lacking cGAS protein.

In-solution fluorescence resonance energy transfer (FRET) Assay

Labeled Bu-DNA or C-dsDNA (2nM Cy3-Cy5 donor–acceptor pair, Cy3 on 5' of 88 bp DNA forward and Cy5 on 5' of Bu-DNA reverse or C-dsDNA reverse as described above, see fig. S4D for illustration) were incubated with 100 nM $cGAS^{FL}$ for 5 min in buffer with 25 mM Tris acetate pH 7.4, 125 mM potassium-acetate pH 7.4, 0.5mM TECP, 5 mM $Mg(Ac)_2$ at pH 7.4, and 5%

glycerol. FRET efficiency was recorded for the Cy3-Cy5 FRET pair with a Tecan M1000 plate reader as previously reported *(91)*.

Single molecule fluorescence resonance energy transfer (smFRET) Assay

Bu-DNA and C-dsDNA oligonucleotide constructs were synthesized and dual-labeled by Integrated DNA Technologies (IDT, Coralville, IA). The donor fluorophore (Cy3) was attached to the 5′ end of one strand, the acceptor (Cy5) was placed on the complementary strand, and a 5′ biotin was introduced to allow surface immobilization via neutravidin. The sequences used (5' to 3') were shown in table S1. Oligonucleotides were annealed by mixing equimolar amounts in annealing buffer (30mM HEPES, pH7.5, 100mM potassium acetate, 1 mM EDTA), heating to 95 °C for 5 min, and then slowly cooling to room temperature over about 2 h on the benchtop. Annealed duplexes were stored at -20 °C and diluted to 1.5 pM immediately before imaging.

For surface preparation and immobilization, quartz microscope slides and coverslips were cleaned by immersion in a freshly prepared basic piranha solution ($H_2O$: $NH_4OH$: $H_2O_2$ in 7:1:1 ratio, heated to ~90 °C for 40 min), followed by thorough rinsing with ultrapure water. The slides were then sequentially sonicated in 1M KOH and methanol. Surfaces were functionalized with aminopropylsilane and polyethylene glycol (PEG) (99 % mPEG-NHS: 1 % biotin-PEG-NHS) to minimize nonspecific adsorption and allow for neutravidin attachment. Flow chambers were assembled using double-sided adhesive spacers, sealed with epoxy and stored at -20 °C. Before sample loading, channels were incubated with 0.2 mg $mL^{-1}$ neutravidin for 5 min, followed by rinsing with imaging buffer (50 mM Tris-HCl pH 7.5, 35 mM KCl, 5 mM $MgCl_2$, 0.07 % IGEPAL, 5 % glycerol). Biotinylated Bu-DNA or C-dsDNA duplexes were then introduced and allowed to bind via biotin-neutravidin interaction for 5 min. Unbound molecules were removed by flushing with the same imaging buffer. For all experiments, cGAS was manually injected into the flow chamber at approximately 20 s after the start of image acquisition.

smFRET data were collected on a prism-type total internal reflection fluorescence (TIRF) microscope built around an inverted Nikon Ti2 frame. Donor excitation was achieved using a 532 nm LED laser (Coherent), and emission was collected through a 60x 1.45 NA oil-immersion objective. Fluorescence was split into donor (582/64 nm) and acceptor (680/42 nm) channels using a 610 nm dichroic mirror and imaged side-by-side on an EMCCD camera (Andor iXon Ultra 897) with 130 ms exposure per frame. Acceptor excitation was achieved using a 637 nm LED laser (Coherent). Movies consisted of 532 nm illumination for >100 sec, followed by 637 nm illumination for 1-2 sec to determine Cy5 photobleaching. The imaging buffer contained an enzymatic oxygen scavenging system composed of 0.1% (w/v) glucose, 100 U $mL^{-1}$ glucose oxidase, and 600 U $mL^{-1}$ catalase, supplemented with 1 mM Trolox to suppress photobleaching and blinking. All experiments were carried out at room temperature (20-22 °C).

Movies were processed using custom Python scripts. Donor-channel fluorescence spots were identified by intensity thresholding and centroid localization, and corresponding acceptor-channel positions were mapped using transformation matrix of stacked frames. Background was subtracted locally for each molecule, and donor and acceptor intensity trajectories were extracted. Corrections for donor bleed-through ($\alpha$) and direct acceptor excitation ($\delta$) were determined from single-color controls and applied. All traces with acceptor (Cy5) photobleaching were excluded from analysis. For each pair of donor/acceptor intensity trajectories, FRET trajectories were calculated as (acceptor/(donor + acceptor)). FRET traces were then plotted, visually inspected, and classified as static, or dynamic based on clear anti-correlated donor/acceptor transitions.

Fluorescence-anisotropy (FA) binding assays

Serially diluted $cGAS^{FL}$ (from 2000 nM to 1.953 nM) or $cGAS^{CAT}$ (from 6000 nM to 7.8125 nM) protein was added to the fluorescein-amidite (FAM)-labeled 88 bp Bu-DNA or C-dsDNA (5 nM final) in buffer (25 mM Tris-acetate (Ac) pH 7.4, 180 mM KAc, 1 mM TCEP, 5 mM $Mg(Ac)_2$, pH 7.4, and 5% glycerol) at 25 ± 2°C. The FA signal was recorded with a Tecan M1000 plate reader as previously reported *(82, 91, 92)*. Changes in FA were plotted as a function of cGAS concentration and fit to the Hill equation. The lines were fit to a standard binding isotherm to obtain the dissociation constant (Kd). For competition-based experiments, unlabeled C-dsDNA or Bu-DNA was titrated against 5 nM of FAM-dsDNA 72 bp and $cGAS^{FL}$ (500 nM). The fraction of bound FAM-dsDNA was quantified and plotted as a function of concentrations of competitor DNAs. Data were fitted to a competition binding model equation: $(1/\ (1+(\text{dsDNA competitor})/IC_{50})^{\text{Hill constant}})$ to derive the $IC_{50}$ values.

Cryo-EM Sample Preparation and Data Acquisition

The cGAS-Bu-DNA88 complex was prepared using size exclusion chromatography (SEC). Purified $cGAS^{FL}$ (Hisx6-MBP tag-TEV protease recognition site-mouse cGAS, 100 µL 37.38mg/mL) was mixed with Bu-DNA88 (24.92 µL of 500 µM) in 875 µL SEC buffer (20 mM Tris at pH 7.5, 150 mM NaCl, and 0.5 mM TCEP) and incubated on ice for 30 minutes before loading on a gel filtration column (HiLoadTM 16/600 Superdex™ 200 pg, Cytiva) pre-equilibrated with SEC buffer.

The peak fraction was used for Cryo-EM sample preparation. Specifically, 4 µL of the complex sample (measured at 1.2 mg/mL of cGAS protein concentration by Bradford protein assay) was applied to Quantifoil® Holey Carbon R 2/2 300 Mesh Au grid (Electron Microscopy Sciences, Q3100AR2) and glow discharged with a PELCO easiGlow (TED PELLA, Inc.) with settings as follows: Plasma Current 15 mA, Process Timer 60s, Working Vacuum 0.37 mBar, Preprocess

Hold Timer 10s, and negative polarity. After 30s wait time, the sample-loaded grid was blotted (blot force 2 and blot time 2s) and plunged frozen in liquid ethane using a Vitrobot Mark IV (Thermo Fisher) at 4°C in 100% humidity. The grid was imaged on a Titan Krios equipped with a Gatan K3 camera and an energy filter operated with a 10-eV width slit (Johns Hopkins Beckman Center for cryoEM). Movies of 40 frames were collected in counting mode with a pixel size of 0.93 Å/pixel and a total dose of 40 $e^-/Å^2$. Data were collected automatically by EPU (Thermo Fisher) using a beam-image shift data collection method with a defocus range of 1.2 to 1.8 µm. A total of 9,927 movies were collected from one session.

Cryo-EM Data Processing

All Cryo-EM data were processed on CryoSPARC (v4.3.1). Movies (EER format) were imported into CryoSPARC for patch motion correction and patch CTF (contrast transfer function) estimation with default settings. A total of 2,969,717 particles were auto-picked using Blob Picker and extracted with a box size of 512 pixels. After two rounds of 2D classifications, 665,277 particles for the cGAS dimer complex were selected for heterogeneous refinement. Best class particles (216,848) were used for homogeneous refinement (3.05Å). Next, the map was flipped along the Z-axis (to flip the handedness) and refined with local CTF refinement (2.89Å). The resulting map was further refined using homogeneous refinement in combination with C2 symmetry to generate the final Cryo-EM density map (2.75Å). For the cGAS tetramer complex, 63,157 particles were selected for heterogeneous refinement. The best class was further refined using non-uniform refinement (3.85Å) and CTF refinement (3.56Å). Finally, the map was refined with non-uniform refinement with C2 symmetry applied (3.48Å). The data processing workflow is in fig. S3A.

Model Building

A crystal structure for the mouse cGAS catalytic domain-dsDNA18 complex (PDB ID 7UUX) was fitted into the final density map of the cGAS dimer complex using UCSF Chimera (v.1.15). The fitted model was then used as an initial model for DNA bubble building in Coot (v.0.9.6). A crystal structure for the mouse cGAS catalytic domain-dsDNA39 complex (PDB ID 5N6I) was fitted into the final density map of the cGAS tetramer complex using UCSF Chimera (v.1.15). The fitted model was then used as an initial model for DNA building in Coot (v.0.9.6). The final models for both structures were refined with real-space refinement in Phenix (v.1.20.1). Information for the deposited map and modeling is summarized in table S2.

Statistical analysis

All statistical analysis was performed in GraphPad Prism 10.4.0 software. Statistical significance was determined by a two-tailed Student's t-test, one-way ANOVA or two-way ANOVA with multiple comparisons and data with error bars were depicted as the average ± SEM or average ± SD. A p value less than 0.05 is considered as significant. *$p<0.05$; **$p<0.01$; ***$p<0.001$.

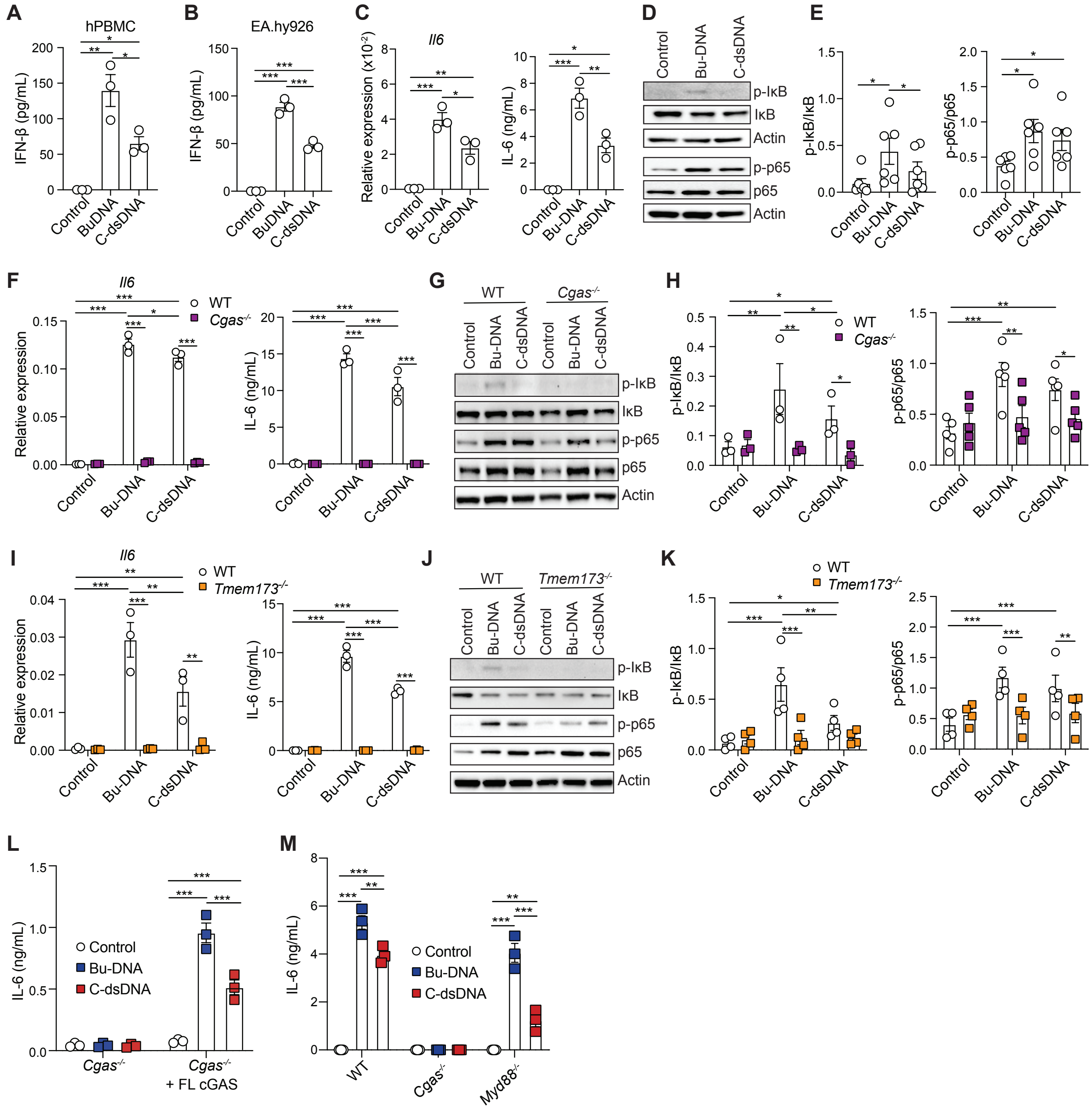

**Fig. S1. Bu-DNA-induced IL-6 responses are cGAS- and STING-dependent**

(**A-B**) Human CD14$^{+}$ monocytes isolated from hPBMC (A) and human endothelial cell line EA.hy926 (B) were transfected with Bu-DNA or C-dsDNA for 18 hours and analyzed for IFN-β

(**C-K**) Mouse BMDMs transfected with Bu-DNA, C-dsDNA for 6 or 18 hours, or left untreated were analyzed for mRNA (C, F and I, left) and protein (C, F and I, right) levels of IL-6. Western blots of indicated targets were shown in (D), (G) and (J), and densitometry in (E), (H) and (K). WT BMDMs were used for C-E, WT and *Cgas*$^{-/-}$ BMDMs for F-H and WT and *Tmem173*$^{-/-}$ BMDMs for I-K.

(**L**) IL-6 levels in *Cgas*$^{-/-}$ iBMDM, unmodified or reconstituted with of cGAS$^{FL}$ transfected with Bu-DNA, C-dsDNA for 18 hours, or left untreated.

(**M**) IL-6 levels in WT, *Cgas*$^{-/-}$ or *Myd88*$^{-/-}$ BMDMs transfected with Bu-DNA, C-dsDNA for 18 hours, or left untreated.

At least three independent experiments were performed with representative results shown. Each dot in (C, F, and I), represents one biological control in one independent experiment. Error bars depict the average ± SEM. Comparisons are made within the same cells for (J and K). *p<0.05; **p<0.01; ***p<0.001.

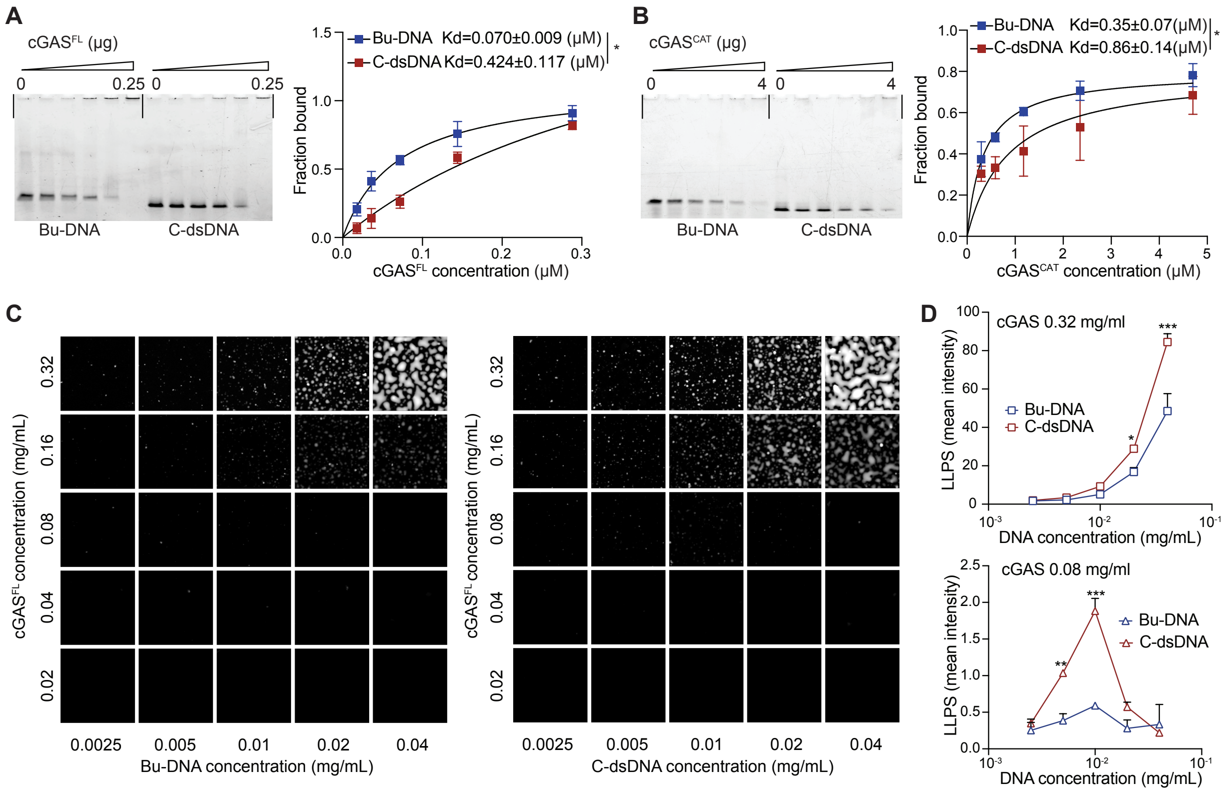

**Fig. S2. cGAS preferentially binds Bu-DNA without further promoting LLPS condensate formation.**

(**A-B**) EMSA experiments assessing the binding of cGAS$^{FL}$ (A) or cGAS$^{CAT}$ (B) to Bu-DNA or C-dsDNA. Recombinant cGAS$^{FL}$ (0 to 0.25 μg, A) or cGAS$^{CAT}$ protein (0 to 4 μg, B) was incubated with 20 ng Bu-DNA or C-dsDNA. A representative gel of four independent experiments is shown on the left. Kd values were determined by fraction bound against protein concentrations (μM) (n=4, right). Error bars depict the mean ± SEM. Statistical significance is calculated by two-tailed unpaired t-test. *p<0.05.

(**C-D**) Confocal images of LLPS of cGAS$^{FL}$-FITC incubated with 60 bp HSV derived Bu-DNA (C, left) or C-dsDNA (C, right) at the indicated concentrations. Mean intensity of images at specific protein concentrations (top, 0.32 mg/mL; bottom, 0.08 mg/mL) were plotted against input DNA concentrations in D. Error bars depict the mean ± SEM. Statistical significance is calculated by two-way ANOVA with multiple comparisons. *p<0.05, ***p<0.001.

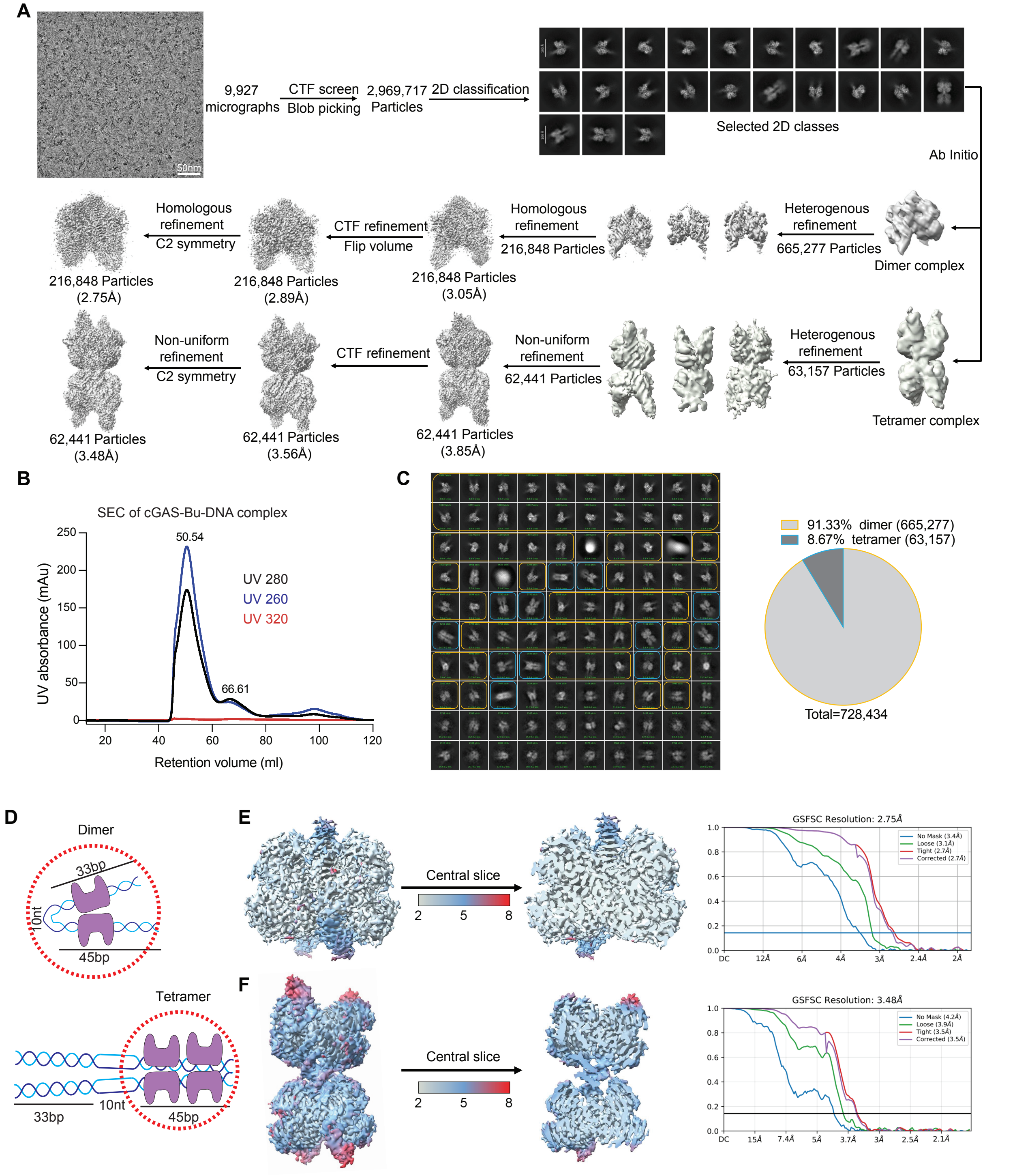

**Fig. S3. Cryo-EM sample preparation of cGAS-Bu-DNA complexes and data processing.**
(**A**) Workflow of cGAS-Bu-DNA complex using CryoSparc.
(**B**) SEC (Superdex 200pg, 16/600) of cGAS and Bu-DNA complex. UV absorbances at 260, 280, and 320 nm were recorded via FPLC detection.
(**C**) 2D classification and particle population analysis of cGAS dimer vs. tetramer complexes. Initial 2D classification into 100 classes were performed to remove low-quality particles, yielding

849,783 particles across 14 classes. These particles underwent a second round of 2D classification into 100 classes. Particle numbers for cGAS dimer complexes (58 classes; 665,262 particles, in yellow boxes) and cGAS tetramer complexes (12 classes; 63,157 particles, in blue box) were quantified and plotted as the percentages of dimer vs tetramer complexes in the pie chart (right).
(**D**) Schematic illustration of cGAS dimer and tetramer complexes bound to Bu-DNA, highlighted in dotted red circle. DNA duplex is depicted in light and dark blue and cGAS in purple.
(**E-F**) Cryo-EM density maps of the cGAS-Bu-DNA dimer (E, left) and tetramer structure (F, left) colored by local resolution (Å). Gold Standard Fourier Shell Correlation (GSFSC) curves for corresponding reconstructions are shown on the right (E and F) with the threshold FSC = 0.143.

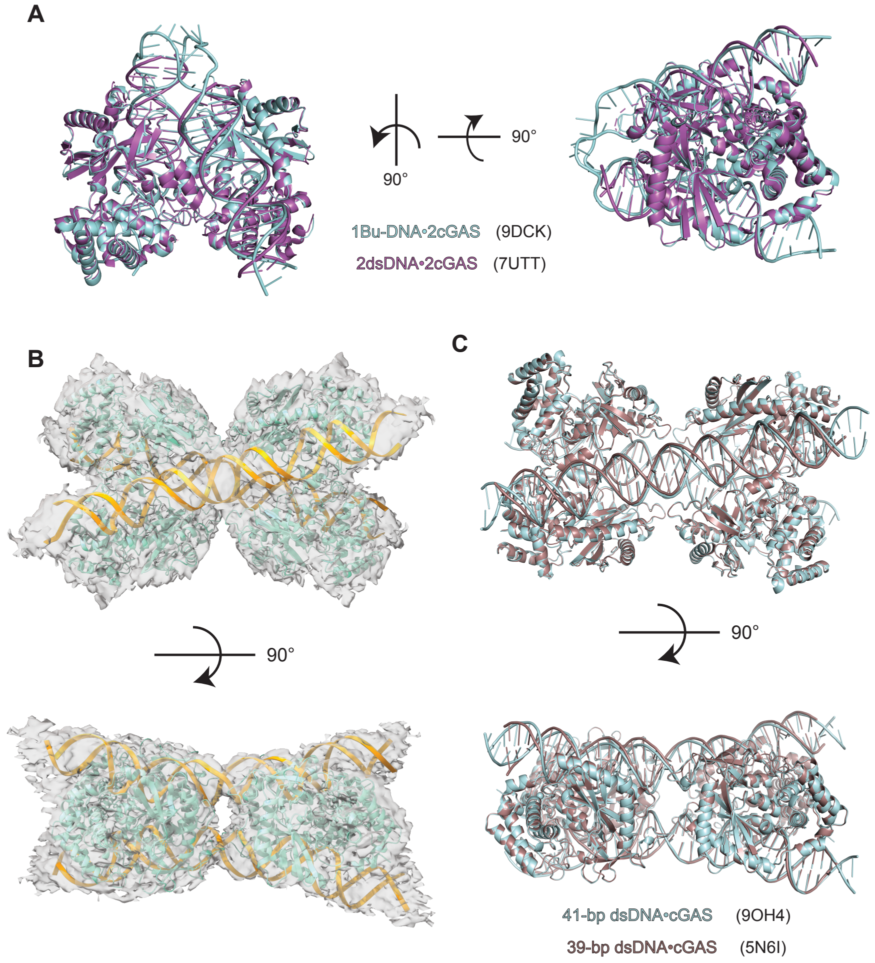

**Fig. S4. Comparison of cGAS-Bu-DNA dimer or tetramer structures with previously reported cGAS-dsDNA complexes**

(**A**) Structural superposition of the cGAS dimer in complex with Bu-DNA from this study (PDB 9DCK, teal) with the previously determined 2:2 cGAS dimer structure (PDB 7UTT, purple).

(**B**) Atomic models of cGAS-Bu-DNA tetramer complex fitted in 3D cryo-EM density maps. cGAS is shown in teal and Bu-DNA is shown in yellow.

(**C**) Structural superposition of the cGAS-Bu-DNA tetramer from this study (PDB 9OH4, teal) with previously determined cGAS tetramer structure (*9*) (PDB 5N6I, brown).

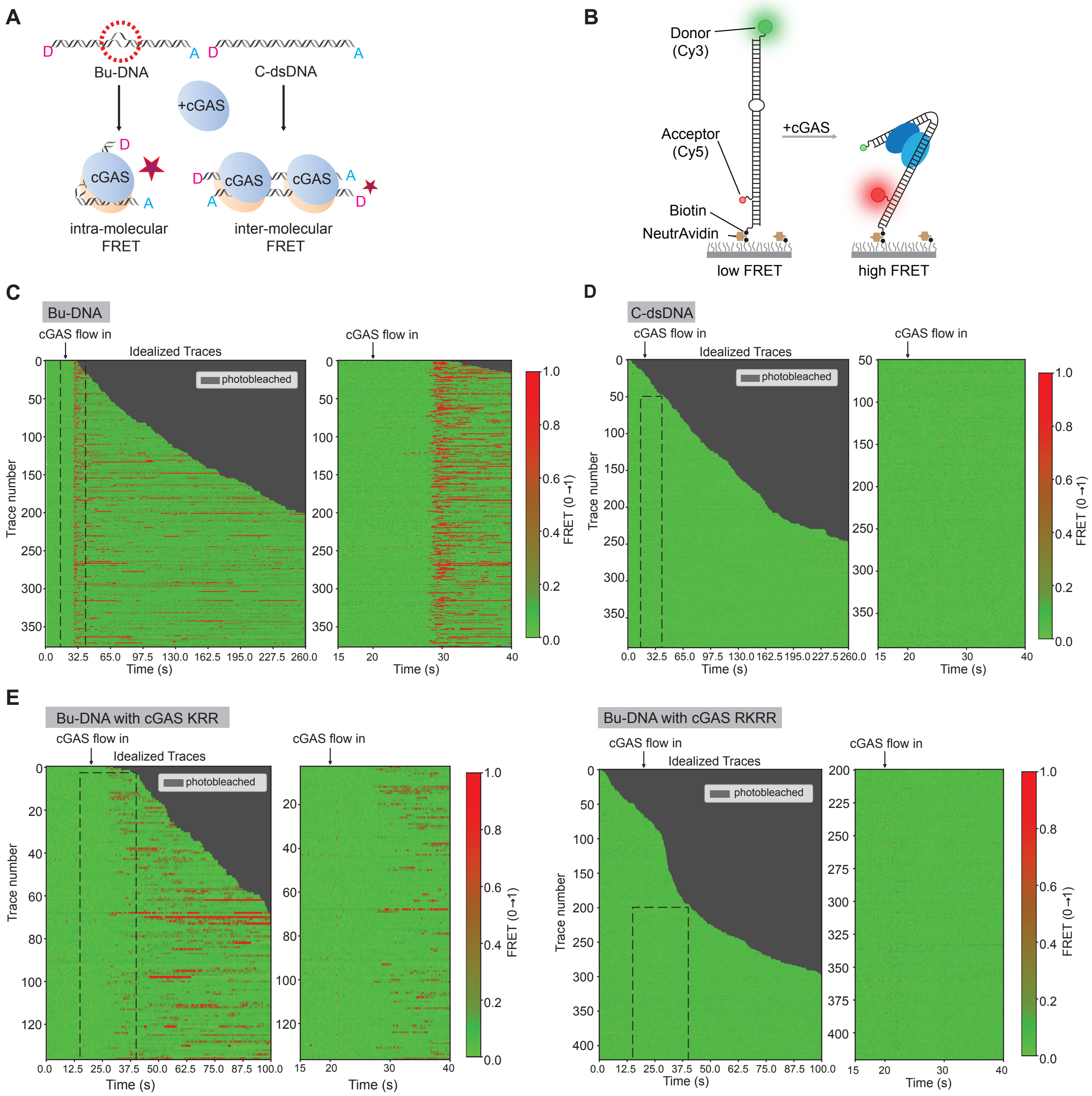

**Fig. S5. Bulk and smFRET assays of cGAS with Bu-DNA vs C-dsDNA**

(**A**) Schematic of the bulk FRET assay. cGAS binding induces energy transfer from donor (D) to acceptor (A) on labeled DNAs (Bu-DNA or C-dsDNA). Bu-DNA exhibits intra-molecular FRET within a single molecule, whereas C-dsDNA only supports inter-molecular FRET between two DNA molecules.

(**B**) Schematic of smFRET assay. Bu-DNA and C-dsDNA were labeled with donor fluorophore Cy3, acceptor fluorophore Cy5, and a biotin to allow surface immobilization via neutravidin. smFRET images were acquired as cGAS was flowed into the imaging chamber.

(**C-E**) Population rastergrams of idealized smFRET trajectories, with each row representing one molecule's FRET efficiency over time. "cGAS flow in" marks the time point of cGAS addition. The right panels are magnified views of the time window indicated by dashed boxes in the left panels. WT cGAS with Bu-DNA (C) or C-dsDNA (D); Bu-DNA with cGAS mutants (E).

**Table S1. DNAs and primers**

| **DNAs** | |
|---|---|
| 88 bp Bu-DNA F: | 5'- GTC GAC GGA ATT CTG AAG TAG GAT TAA TAG TAG TTT TTT TTT TCA CAG AGA AGA ACA TTT GAC CCG GGT AAA GCT AAT AAC AAG TAA T -3' |
| 88 bp Bu-DNA R: | 5'- ATT ACT TGT TAT TAG CTT TAC CCG GGT CAA ATG TTC TTC TCT GTG CCC CCC CCC CCT ACT ATT AAT CCT ACT TCA GAA TTC CGT CGA C -3' |
| 88 bp C-dsDNA R: | 5'- ATT ACT TGT TAT TAG CTT TAC CCG GGT CAA ATG TTC TTC TCT GTG AAA AAA AAA ACT ACT ATT AAT CCT ACT TCA GAA TTC CGT CGA C-3' |
| Cy3-Bu-DNA F: | 5'Cy3- GTC GAC GGA ATT CTG AAG TAG GAT TAA TAG TAG TTT TTT TTT TCA CAG AGA AGA ACA TTT GAC CCG GGT AAA GCT AAT AAC AAG TAA T -3' |
| Cy5-Bu-DNA R: | 5'Cy5- ATT ACT TGT TAT TAG CTT TAC CCG GGT CAA ATG TTC TTC TCT GTG CCC CCC CCC CCT ACT ATT AAT CCT ACT TCA GAA TTC CGT CGA C -3' |
| Cy5-C-dsDNA R: | 5'Cy5- ATT ACT TGT TAT TAG CTT TAC CCG GGT CAA ATG TTC TTC TCT GTG AAA AAA AAA ACT ACT ATT AAT CCT ACT TCA GAA TTC CGT CGA C-3' |
| smFRET Cy3 oligo: | 5′Cy3-GT CGA CGG AAT TCT GAA GTA GGA TTA ATA GTA GTT TTT TTT TTC ACA GAG AAG AAC ATT TGA CCC GGG TAA AGC TAA TAA CAA GTA ATA GA CAG CTG CACG ACG TTG CG -3′ |
| smFRET Cy5 oligo Bu-DNA: | 5’Cy5- AT TAC TTG TTA TTA GCT TTA CCC GGG TCA AAT GTT CTT CTC TGT GCC CCC CCC CCC TAC TAT TAA TCC TAC TTC AGA ATT CCG TCG AC -3’ |
| smFRET Cy5 oligo C-dsDNA: | 5’Cy5- AT TAC TTG TTA TTA GCT TTA CCC GGG TCA AAT GTT CTT CTC TGT G**AA AAA AAA AA**C TAC TAT TAA TCC TAC TTC AGA ATT CCG TCG AC -3’ |
| smFRET biotin handle strand: | 5’Biotin- A ACG CAA CGT CGT CAG CTG TCT AA GCA GCT GTC TAT TAC TTG TTA T -3’ |
| 288bp Bu-DNA F: | 5’-AAA CGA CGG CCA GTG AAA TTT GGT ACC TGA GCA GTT CCC AGC TTG ACT TCG TCC TCA CTC TCT TCC TCT AGC GCT ATAT GCG TTG ATG GAC CAG GAC CAG GTC GAC GGA ATT CTG AAG TAG GAT TAA TGA GAA ACC AAC CAA CCA ACC AAC CAA CAC ATT TGA CCC GGG TAA AGC TAA TAA CAA GTA ACT GGT CCT GGT CCA TCA ACG CAT ATA GCG CTA GAG GAA GAG AGT GAG GAC GAA GTC AAG CTG GGA ACT GCT CAG GTA CCA AAT TTC ACT GGC CGT CGT TT-3’ |

| | |
|---|---|
| 288bp Bu-DNA R: | 5'- AAA CGA CGG CCA GTG AAA TTT GGT ACC TGA GCA GTT CCC AGC TTG ACT TCG TCC TCA CTC TCT TCC TCT AGC GCT ATA TGC GTT GAT GGA CCA GGA CCA GTT ACT TGT TAT TAG CTT TAC CCG GGT CAA ATG TTA ACC CTA ACC CTA ACC CTA ATT TCT CAT TAA TCC TAC TTC AGA ATT CCG TCG ACC TGG TCC TGG TCC ATC AAC GCA TAT AGC GCT AGA GGA AGA GAG TGA GGA CGA AGT CAA GCT GGG AAC TGC TCA GGT ACC AAA TTT CAC TGG CCG TCG TTT -3' |
| 288bp C-dsDNA R: | R: 5'-AAA CGA CGG CCA GTG AAA TTT GGT ACC TGA GCA GTT CCC AGC TTG ACT TCG TCC TCA CTC TCT TCC TCT AGC GCT ATA TGC GTT GAT GGA CCA GGA CCA GTT ACT TGT TAT TAG CTT TAC CCG GGT CAA ATG TGT TGG TTG GTT GGT TGG TTG GTT TCT CAT TAA TCC TAC TTC AGA ATT CCG TCG ACC TGG TCC TGG TCC ATC AAC GCA TAT AGC GCT AGA GGA AGA GAG TGA GGA CGA AGT CAA GCT GGG AAC TGC TCA GGT ACC AAA TTT CAC TGG CCG TCG TTT-3' |
| 170bp Cir-BuDNA F: | 5'Phos- AGG TCG CCG CCC CGT AAC AC CCT TGA CGA GTC CAT GGT TTA AAG TGA CCG GCA GCA TTA CTT GTT ATT AGC TTT ACC CGG GTC AAA TGT TAA CCC TAA CCC TAA CCC TAA TTC TCA TTA ATC CTA CTT CAG AAT TCC GTC GAC CTG GTC CTG GTA GTC TGC GT ACCCGCG -3' |
| 170 bp Cir-Bu-DNA R: | 5'Phos- G GGC GGC GAC CTC GCG GGT AC GCA GAC TAC CAG GAC CAG GTC GAC GGA ATT CTG AAG TAG GAT TAA TGA GAA ACC AAC CAG CCA CCG CCA ACA ACA TTT GAC CCG GGT AAA GCT AAT AAC AAG TAA TGC TGC CGG TCA CTT TAA ACC ATG GAC TCG TCA AGG GT GTTACG -3'. |
| 170bp Cir-C-dsDNA F: | 5'Phos- G GGC GGC GAC CTC GCG GGT AC GCA GAC TAC CAG GAC CAG GTC GAC GGA ATT CTG AAG TAG GAT TAA TGA GAA TTA GGG TTA GGG TTA GGG TTA ACA TTT GAC CCG GGT AAA GCT AAT AAC AAG TAA TGC TGC CGG TCA CTT TAA ACC ATG GAC TCG TCA AGG GT GTTACG -3' |
| **RT-PCR primers** | |
| *Ifnb* F: | 5'- CCA GCT CCA AGA AAG GAC GA -3' |
| *Ifnb* R: | 5'- CGC CCT GTA GGT GAG GTT GAT -3' |
| *Il6* F: | 5'- CCA GAG TCC TTC AGA GAG ATA CA -3' |
| *Il6* R: | 5'- CCT TCT GTG ACT CCA GCT TAT C -3' |
| *Dloop* F: | 5'- AATCTACCATCCTCCGTGAAACC -3' |

| | |
|---|---|
| *Dloop* R: | 5’- TCAGTTTAGCTACCCCCAAGTTTAA -3’ |
| *Cytb* F: | 5’- GCTTTCCACTTCATCTTACCATTTA -3’ |
| *Cytb* R: | 5’- TGTTGGGTTGTTTGATCCTG -3’ |
| *Tert* F: | 5’- CTAGCTCATGTGTCAAGACCCTCTT -3’ |
| *Tert* R: | 5’- GCCAGCACGTTTCTCTCGTT -3’ |
| *Actb* F: | 5’- CAT TGC TGA CAG GAT GCA GAA GG -3’ |
| *Actb* R: | 5’- TGC TGG AAG GTG GAC AGT GAG G -3’ |
| *Gapdh* F: | 5’- CAT CAC TGC CAC CCA GAA GAC TG -3’ |
| *Gapdh* R: | 5’- ATG CCA GTG AGC TTC CCG TTC AG -3’ |
| *B2m* F: | 5’- ACA GTT CCA CCC GCC TCA CAT T -3’ |
| *B2m* R: | 5’- TAG AAA GAC CAG TCC TTG CTG AAG -3’ |

**Table S2. Cryo-EM data collection, refinement, and validation statistics**

| | cGAS dimer-BuDNA88 (EMDB-46754) (PDB 9DCK) | cGAS tetramer-BuDNA88 (EMDB-70483) (PDB 9OH4) |
|---|---|---|
| **Data collection and processing** | | |
| Magnification | 130,000x | 130,000x |
| Voltage (kV) | 300 | 300 |
| Electron exposure ($e^{-}/Å^{2}$) | 40 | 40 |
| Defocus range (μm) | -1.2 to -1.8 | -1.2 to -1.8 |
| Pixel size (Å) | 0.93 | 0.93 |
| Symmetry imposed | C2 | C2 |
| Initial particle images (no.) | 2,969,717 | 2,969,717 |
| Final particle images (no.) | 216,848 | 62,441 |
| Map resolution (Å)<br>FSC threshold = 0.143 | 2.75 | 3.48 |
| Map resolution range (Å) | 2.2 – 8.3 | 2.3 – 11.5 |
| **Refinement** | | |
| Initial model used (PDB code) | 7UUX | 5N6I |
| Model resolution (Å)<br>FSC threshold = 0.143 | 2.7 | 3.5 |
| Model resolution range (Å) | 2.7 – 3.1 | 3.4 – 3.8 |
| Map sharpening *B* factor ($Å^{2}$) | -60.2 | -87.5 |
| Model composition<br>Non-hydrogen atoms<br>Protein residues<br>Nucleotide<br>Ligands (Zn) | <br>7,803<br>718<br>92<br>2 | <br>29,047<br>1,248<br>164<br>4 |
| *B* factors ($Å^{2}$)<br>(min/max/mean)<br>Protein<br>Nucleotide<br>Ligand | <br>2.67/7.46/3.38<br>2.67/100.32/38.65<br>30.92/39.36/35.14 | <br>31.42/110.29/63.39<br>48.27/151.13/92.18<br>52.91/68.68/62.03 |
| R.m.s. deviations<br>Bond lengths (Å)<br>Bond angles (°) | <br>0.004<br>0.639 | <br>0.003<br>0.652 |
| Validation<br>MolProbity score<br>Clashscore<br>Poor rotamers (%) | <br>1.79<br>8.06<br>0.30 | <br>2.49<br>10.31<br>2.73 |
| Ramachandran plot | | |

| | | |
|---|---|---|
| Favored (%) | 94.96 | 91.20 |
| Allowed (%) | 5.04 | 8.66 |
| Disallowed (%) | 0.00 | 0.14 |

**Movie S1. Bu-DNA and cGAS**

**Movie S2. smFRET of Bu-DNA and cGAS**

**Movie S3. smFRET of C-dsDNA and cGAS**

**Movie S4. smFRET of KRR cGAS and Bu-DNA**

**Movie S5. smFRET of RKRR cGAS and Bu-DNA**
